# Multiple roles for the ESCRT machinery in maintaining plasma membrane homeostasis

**DOI:** 10.1101/2020.02.25.964452

**Authors:** Oliver Schmidt, Yannick Weyer, Simon Sprenger, Michael A. Widerin, Sebastian Eising, Verena Baumann, Mihaela Angelova, Robbie Loewith, Christopher J. Stefan, Michael W. Hess, Florian Fröhlich, David Teis

## Abstract

The endosomal sorting complexes required for transport (ESCRT) execute evolutionary conserved membrane remodeling processes. Here we used budding yeast to explore how the ESCRT machinery contributes to plasma membrane (PM) homeostasis. In response to reduced membrane tension and inhibition of the target of rapamycin complex 2 (TORC2), ESCRT-III/Vps4 assemblies form at the PM and help to maintain membrane integrity. Conversely, the growth of ESCRT mutants strongly depends on TORC2-mediated homeostatic regulation of sphingolipid (SL) metabolism. This is caused by calcineurin phosphatase activity which causes Orm2 to accumulate at the endoplasmic reticulum (ER) in ESCRT mutants. Orm2 is a repressor of SL biosynthesis and its accumulation provokes increased membrane stress. This necessitates TORC2 signaling through its downstream kinase Ypk1 to control Orm2 protein levels and prevent a detrimental imbalance of SL metabolism. Our findings reveal new aspects of antagonistic calcineurin/TORC2 signaling for the regulation of SL biosynthesis and the maintenance of PM homeostasis, and suggest that the ESCRT machinery contributes directly and indirectly to these processes.

## Introduction

Eukaryotic cells harmonize lipid homeostasis and protein homeostasis (proteostasis) to maintain the integrity of their membranes. This requires the balanced synthesis of proteins and lipids and their selective degradation. How these processes are coordinated is only partially understood.

Selective membrane protein degradation is mediated through several ubiquitin-dependent degradation pathways. In the ER, the multi-subunit transmembrane ubiquitin ligases of the ER-associated degradation (ERAD) machinery ubiquitinate membrane proteins for retro-translocation into the cytoplasm and subsequent proteasomal degradation (1, 2). ERAD can also degrade functional proteins in a regulated manner and thereby control membrane homeostasis. An important regulated ERAD substrate is the 3-hydroxy-3-methylglutaryl coenzyme A reductase (HMGR), a key enzyme in sterol biosynthesis (3, 4). HMGR degradation is part of a rheostat that is critical for sterol homeostasis in yeast and humans. Also, squalene monooxygenase is subject to sterol-regulated ERAD (5, 6). When the folding and degradation capacity of the ER and ERAD are overwhelmed, mis-folded proteins accumulate and induce a cellular stress response pathway, called the unfolded protein response (UPR). The UPR implements adaptive transcriptional programs that resolve ER stress by a decrease in overall protein synthesis and a specific up-regulation of hundreds of genes including lipid biosynthesis, ER chaperons and ERAD machineries (7). Consistently, simultaneous loss of ERAD and UPR decreases viability of yeast cells (8).

Proteins in post-ER organelles are degraded by different degradation pathways. We recently described the endosome and Golgi-associated degradation (EGAD) pathway (9). EGAD is mediated by the defective in sterol-responsive element binding protein (SREBP) cleavage (Dsc) complex, a multi-subunit transmembrane ubiquitin ligase reminiscent of the complexes operating in ERAD (10). During EGAD, the Dsc complex ubiquitinates membrane proteins at the Golgi and endosomes and thereby initiates their extraction into the cytosol by the AAA (ATPase associated with diverse cellular activities) protein Cdc48, which leads to their proteasomal degradation (9). The Dsc complex also ubiquitinates a distinct set of substrates for lysosomal degradation via the ESCRT machinery (discussed below) (11–14). The EGAD and ESCRT machineries show a strong negative genetic interaction (synthetic growth defect), suggesting that they have non-redundant functions and non-overlapping substrates (9, 14).

A substrate that is exclusively degraded via the EGAD pathway is the transmembrane protein Orm2 (9). Orm2 and its paralog Orm1 are critical components of the multi-subunit SPOTS complex in the ER. The SPOTS complex consists of Orm1/2, the catalytic subunits of the serine-palmitoyl-coenzyme A transferase (SPT, Lcb1 and Lcb2) and of the regulatory factors Tsc3 and Sac1 (15, 16). The SPT mediates sphingoid long chain base (LCB) synthesis and is rate limiting for sphingolipid metabolism. The Orm1/2 proteins repress SPT activity and ensure that the SPT only produces LCBs that are subsequently consumed in SL biosynthesis. Thereby Orm1/2 prevent the toxic accumulation of LCBs. The function of Orm1/2 is evolutionary conserved. Loss of ORMDL1-3, their mammalian orthologues, results in neurotoxicity in mice (17), whereas increased expression of ORMDL3 causes imbalances in sphingolipid metabolism in humans, and is associated with childhood asthma, Crohn’s disease, ulcerative colitis and diabetes (18–21). In yeast, the de-repression of SPT activity is mediated by phosphorylation of the Orm1/2 proteins in their N-terminal cytosolic domain by the AGC-family serine-threonine protein kinase Ypk1 (22). Phosphorylated Orm2 is exported from the ER to Golgi and endosomes, where the protein is degraded by the EGAD machinery (9).

Ypk1, the yeast orthologue of the mammalian serum- and glucocorticoid-induced protein kinase SGK1, is activated through phosphorylation by two upstream kinases. Phosphorylation of the activation loop by phosphoinositide-dependent kinase (Pkh1/2 in yeast) is essential for activity, but full activation requires phosphorylation from the target of rapamycin complex 2 (TORC2) (23). In yeast, TORC2 is an essential, PM-localized multi-subunit protein kinase whose activity responds to membrane stress conditions such as changes in PM tension (24, 25) and sphingolipid levels (26). How TORC2 senses membrane stress is largely unknown, but its regulation involves the dynamic partitioning of TORC2 subunits into membrane subdomains which can either promote (24, 27) or inhibit (25) activity. In addition TORC2 can be regulated by association with Rab5-family proteins (28). Activation of Ypk1 by TORC2 requires its regulated recruitment to the plasma membrane by the TORC2 accessory proteins Slm1/2 (24, 29) and the TORC2 subunit Avo1 (30). The phosphorylation of the Orm1/2 proteins constitutes the essential function of the TORC2-Ypk1 signaling axis (22, 31). Further Ypk1-regulated processes include the stimulation of ceramide synthesis (32), aminophospholipid transport (33), osmoregulation (34, 35) and endocytosis/actin cytoskeleton dynamics (31, 36–38), all impinging in one way or another on membrane homeostasis. Ypk1 has a paralogue, Ypk2, which shares its activation by Pkh1/2 and TORC2. Ypk1 and Ypk2 are believed to share a largely overlapping substrate spectrum. Since the simultaneous deletion of both kinases is lethal, Ypk1 and Ypk2 at least partially compensate for each other’s function (39). However, only Ypk1 activity is required for normal growth of yeast cells.

The degradation of most membrane proteins of the PM, the Golgi or endo- and lysosomal membranes occurs inside lysosomes (vacuoles in yeast) and depends on the ESCRT machinery (40–43). Early ESCRT complexes (ESCRT-0, -I, -II) probably cluster and sequester ubiquitinated cargo proteins on the endosome surface and initiate the formation of the filamentous ESCRT-III complex. Dynamic remodeling of the ESCRT-III complex by the AAA ATPase Vps4 and its partner proteins drives the budding of small cargo-laden vesicles into the lumen of the endosome (44). This process generates multivesicular bodies (MVB), which subsequently fuse with the lysosome/vacuole. Some cargo proteins are also budded by the ESCRT machinery directly from the lysosomal surface into the lumen in response to TORC1 inactivation (45–47). Loss of function of the ESCRT machinery leads to the formation of aberrant, peri-vacuolar endosomal structures called class E compartments, where substrates of the MVB pathway accumulate (48). The ESCRT machinery is additionally involved in many other cellular membrane remodeling events, including abscission of the midbody at the end of cytokinesis, the generation of exosomes, or budding of enveloped viruses (e.g. HIV) (for a recent review on ESCRT functions, see (41, 49)). The ESCRT machinery also repairs ruptured lysosomal, nuclear and plasma membranes (50–58).

While regulatory roles of ERAD and EGAD in lipid metabolism have been described, it is less clear how the ESCRT machinery contributes to membrane homeostasis. Here we show that the ESCRT machinery has an essential and complex role in maintaining PM homeostasis. ESCRT-III/Vps4 assemblies form at the PM in response to reduced PM tension or TORC2 inhibition. Cells without a functional ESCRT machinery are hyper-sensitive to plasma membrane stress conditions and require TORC2-Ypk1 signaling to maintain PM integrity and promote growth and survival. The mutual dependence of the ESCRT machinery and TORC2-Ypk1 signaling is caused by accumulation of Orm2 at the ER. In ESCRT mutants, Orm2 is not efficiently exported from the ER because the phosphatase calcineurin counteracts Orm2 ER export by causing its untimely dephosphorylation. The accumulating Orm2 proteins at the ER trigger elevated TORC2-Ypk1 signaling to maintain homeostasis, and loss of Ypk1 signaling in ESCRT mutants results in a fatal imbalance in SL metabolism with an accumulation of LCBs. Importantly, preventing calcineurin-dependent Orm2 dephosphorylation promoted the degradation of Orm2 by EGAD, and was sufficient to break the dependence of ESCRT mutants on TORC2-Ypk1 signaling. This demonstrates that the accumulation of Orm2 at the ER is a major membrane stressor. Moreover, our results reveal an important link between selective membrane protein degradation systems in post-ER compartments and TORC2-Ypk1 dependent membrane stress signaling.

## Results

### ESCRT mutants depend on TORC2-Ypk1 signaling for cell growth and survival

The evolutionary conserved ESCRT complexes assemble into membrane remodeling machineries that execute fundamental biological processes, including the biogenesis of multivesicular bodies for lysosomal membrane protein degradation and the repair of diverse cellular membranes (41). Given these key roles of the ESCRT machinery, it is surprising that ESCRT mutant cells are viable. This implies that cells can at least in part compensate for the loss of ESCRT function. Understanding how cells achieve this compensation would therefore also provide a better understanding for the role of the ESCRT machinery in cellular homeostatic processes.

To identify the processes that enable the survival of ESCRT-deficient cells, we used the budding yeast, *S. cerevisiae*, as a model system and conducted a genome-wide synthetic genetic interaction screen (9). Gene ontology (GO) term enrichment analysis revealed ‘lipid metabolism’ among the top ranked hits that are required by ESCRT mutants (*vps4*Δ) (Fig. 1A, Table S1). This GO term included genes involved in the metabolism of various lipid classes, but predominantly enzymes mediating non-essential steps of sterol biosynthesis, and genes regulating SL homeostasis. We confirmed the synthetic growth defect of *vps4*Δ cells with four of these genes (*ypk1*Δ, *sac1*Δ, *csg2*Δ, *erg2*Δ) in a different genetic background (the SEY6210 strain) (Fig. S1A). These results suggested that the loss of ESCRT-dependent membrane remodeling rendered cells particularly sensitive to perturbations of membrane homeostasis.

**Figure 1:**
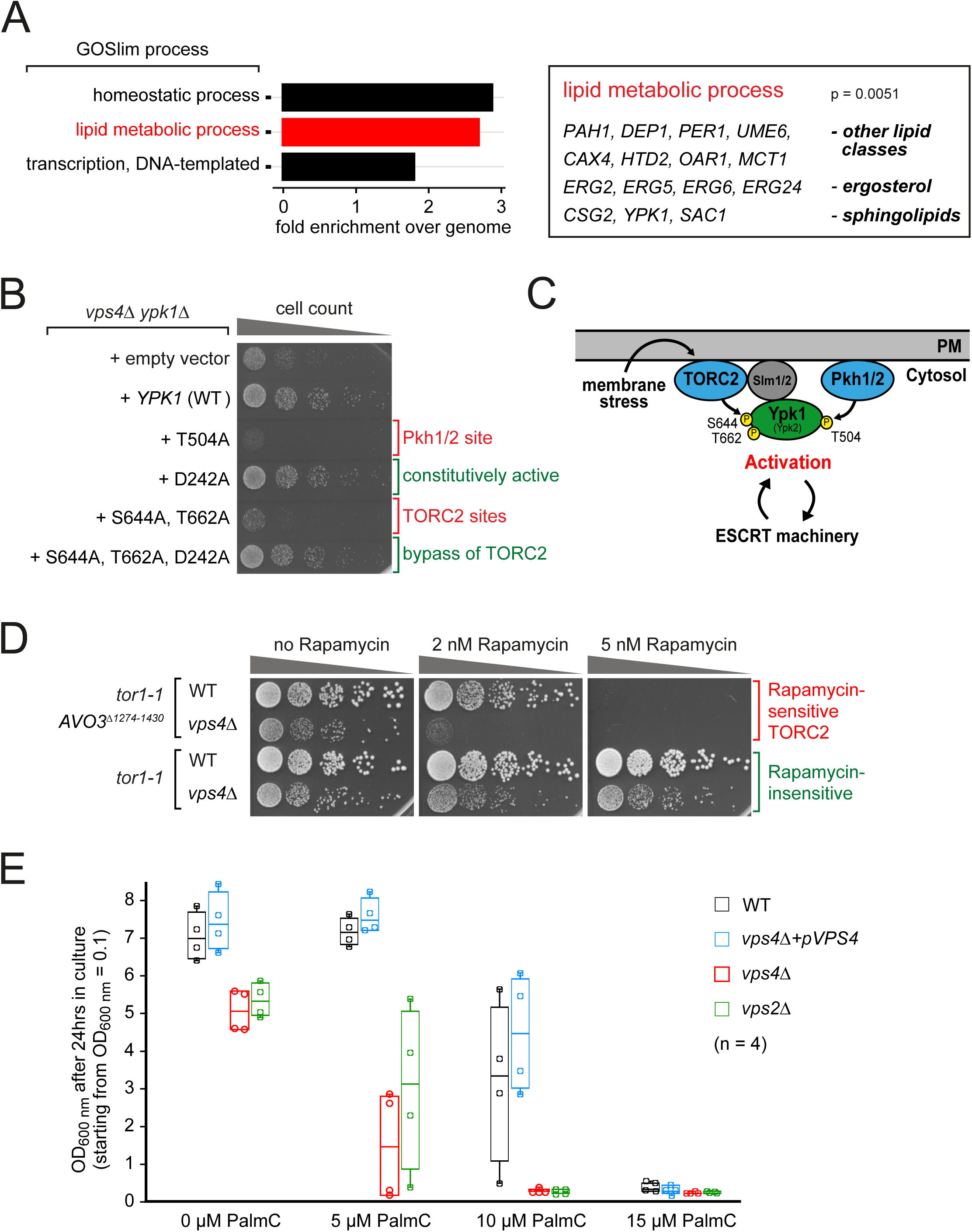
ESCRT mutants depend on lipid metabolism and TORC2-Ypk1 signaling for growth. **(A)** Gene Ontology (GO) analysis for cellular processes of 119 genes that showed negative genetic interactions with *vps4*Δ (9). Only significantly enriched GO terms (p < 0.05) are shown. 15 genes comprised in the GO term ‘lipid metabolic process’ are shown. Full GO analysis in Table S1. **(B)** Equal amounts of *vps4*Δ *ypk1*Δ double mutants expressing the indicated *YPK1* mutant plasmids (or empty vector) in serial dilutions were incubated on auxotrophic selection medium agar plates at 26°C. **(C)** Schematic representation of the activation of the TORC2-Ypk1 signaling pathway in response to membrane stress and its putative interdependence with the ESCRT machinery. PM, plasma membrane; TORC2, target of rapamycin complex 2; Pkh1/2, yeast orthologues of 3-phosphoinositide-dependent kinase 1. **(D)** Equal amounts of the indicated strains in serial dilutions were incubated on auxotrophic auxotrophic selection medium agar plates containing the indicated rapamycin concentrations at 26°C. All WT strains are the respective *vps4*Δ strain re-expressing *VPS4* from plasmid. **(E)** Growth of the indicated strains in presence of palmitoyl-carnitine (PalmC). Cells were inoculated to OD_600nm_ = 0.1 in auxotrophic selection medium containing the indicated PalmC concentrations and grown for 24 hours in 4 independent experiments. The circles indicate the individual measurements. See also Figure S1 and Table S1.

We focused on the role of the protein kinase Ypk1 in ESCRT-deficient cells, because it functions together with TORC2 to maintain membrane homeostasis (22,24,33,36,59,60). *YPK1* deletion (but not deletion of *YPK2*) also caused synthetic growth defects with ESCRT-II (*vps25*Δ) and ESCRT-III mutants (*vps2*Δ), but not with mutants of Chm7 (*chm7*Δ)(Fig. S1B), which recruits the ESCRT machinery to the ER and the inner nuclear membrane (51,52,61). This result suggested that the function of Ypk1 was required to compensate for the loss of ESCRT functions that are independent of Chm7.

The growth of *vps4*Δ *ypk1*Δ double mutants was restored by re-expression of *VPS4* and *YPK1* from plasmids (Fig. S1C), but not by an additional copy of *YPK2* (39, 62). Expression of a Ypk1 mutant (Ypk1 S644A, T662A) that can no longer be activated by TORC2 failed to rescue the growth of *vps4*Δ *ypk1*Δ (Fig. 1B,C) or *vps2*Δ *ypk1*Δ (Fig. S1D) double mutants, similar to the catalytically dead T504A mutant (63). The constitutively active D242A mutant (22) restored normal growth of the double mutants, even when the TORC2 acceptor sites were mutated (Fig. 1B, S1D). Hence, TORC2-dependent Ypk1 activation was required in ESCRT mutants.

To further characterize the relation between TORC2 signaling and the ESCRT machinery, we analyzed the growth of WT and ESCRT mutant cells, which were genetically engineered for the selective inhibition of TORC2 (*tor1-1 AVO3*^Δ*1274-1430*^) with rapamycin (64). At 2 nM rapamycin the growth of WT cells was only mildly affected (64), but the growth of *vps4*Δ mutants was already markedly reduced (Fig. 1D).

TORC2 responds to perturbation of the PM, and acute reduction of PM tension with palmitoyl-carnitine (PalmC) leads to transient TORC2 inactivation (25). Importantly, the growth of WT cells was not affected by low doses of PalmC (5 µM), suggesting that they were capable to maintain homeostasis. Yet, the same PalmC concentration already slowed down the growth of *vps4*Δ or *vps2*Δ cells (Fig. 1E), suggesting that ESCRT function somehow contributed to viability upon PM stress.

Thus, our results imply that TORC2-Ypk1 signaling and the ESCRT machinery have non-redundant functions that together are critical to maintain membrane homeostasis and promote growth.

### ESCRT-III and Vps4 localize to plasma membrane structures upon TORC2 inhibition

TORC2 signaling responds to tensile stress at the PM and in consequence controls sphingolipid synthesis and other processes to maintain PM homeostasis (24,25,65). How the ESCRT machinery contributes to maintaining PM homeostasis was less clear. In human cells, the ESCRT machinery localizes to PM lesions and repairs the ruptured areas, and also prolongs PM integrity during necroptosis (56,66–68). If and how these processes might be correlated to tensile PM stress and TORC2 signaling in budding yeast is unknown.

We therefore analyzed the localization of the major ESCRT-III subunit Snf7 in cells that were treated with 5µM PalmC, a concentration that did not affect the growth of WT cells (Fig. 1E). In untreated cells, Snf7-eGFP was mainly detected in the cytosol and on objects corresponding to endosomes and multivesicular bodies (MVBs) (44) (Fig. 2A, arrowheads). Upon addition of PalmC, we detected the formation of additional Snf7-eGFP foci at or close to the PM (Fig. 2A, arrows). After 2 hours, at least three of these structures were observed in the majority of cells (66%) (Fig. 2A, B). Thus, ESCRT-III assemblies formed at the PM in response to PalmC-induced membrane perturbations.

**Figure 2:**
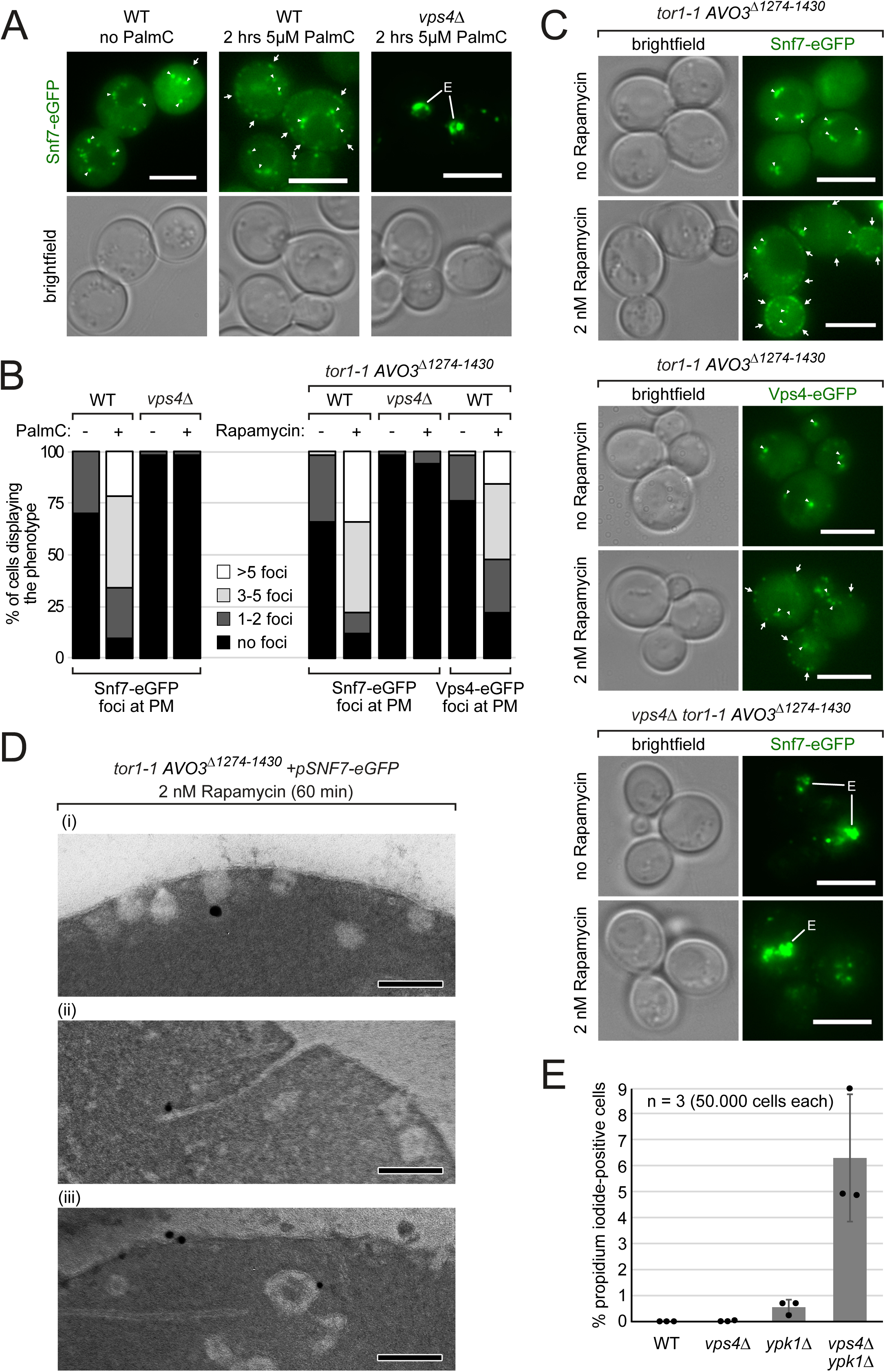
ESCRT-III/Vps4 assemblies form at the plasma membrane. **(A)-(D)** All WT strains are isogenic to the respective *vps4*Δ cells and re-express *VPS4* (or *VPS4-eGFP* where indicated) from a plasmid. **(A)** Epifluorescence and phase contrast microscopy of WT and *vps4*Δ cells co-expressing Snf7-eGFP (green) from a centromeric plasmid and treated with 5 µM palmitoyl-carnitine (PalmC) or vehicle (DMSO) for 2 hours. Arrowheads indicate Snf7 on endosomes/MVBs and arrows indicate Snf7 assemblies at the PM. Scale bars 5µm. **(B)** Quantification of (B) and (C). Sub-cortical eGFP dots in 50 cells of each genotype and condition were counted and grouped into the indicated cohorts. **(C)** Epifluorescence and phase contrast microscopy of the indicated strains co-expressing *SNF7-eGFP* or expressing *VPS4-eGFP* from centromeric plasmids and treated with 2nM rapamycin or vehicle (DMSO) for 2 hours. Arrowheads indicate endosomes/MVBs and arrows indicate Snf7/Vps4 assemblies at the PM. Scale bars 5µm. **(D)** High-pressure freezing, anti-GFP immunogold labeling and transmission electron microscopy of the indicated cells treated with 2nM rapamycin for 60 minutes. Scale bars 200 nm. **(E)** The indicated strains were grown into mid-log phase and stained with propidium iodide for 10 minutes. 50.000 cells were analyzed by flow cytometry. Data are presented as mean ±standard deviation from 3 independent experiments.

Next, we tested if the localization of ESCRT-III to the PM was a consequence of TORC2 inactivation. We determined the localization of Snf7 and of the AAA-ATPase Vps4 upon sublethal inhibition of TORC2 signaling (2 nM rapamycin) (Fig. 1D). Similar to the PalmC-treated cells, Snf7-eGFP and Vps4-eGFP assemblies formed at the PM in many cells (> 75%) after 2 hours of TORC2 inhibition (Fig. 2B, C, arrows). One or two sub-cortical Snf7-eGFP and Vps4-eGFP dots but rarely more were also observed in control WT cells (in 22-34% of cells) (Fig. 2B).

To describe the identity of the Snf7-eGFP positive structures that emerge upon PM stress at the ultrastructural level, we performed cryofixation and immunogold electron microscopy (44). After 60 min of sublethal TORC2 inhibition, we observed numerous vesicular structures at or near the PM (Fig. 2D). PM-associated vesicles likely represented invaginations, possibly reflecting stalled endocytic events, since TORC2 signaling stimulates endocytosis on multiple levels (31,36,65). Snf7-eGFP localized to these invaginations (Fig. 2D (i)), to extended tubular invaginations (ii) as well as to flat areas and vesicular structures in close vicinity to the PM (iii).

The localization of Snf7-eGFP to PM foci in response to PalmC treatment or direct TORC2 inhibition required the activity of Vps4. In *vps4*Δ mutants, Snf7-eGFP remained associated with class E compartments (44) (Fig. 2A, B, C). Of note, the growth of *vps4*Δ cells was already severely affected under these conditions (Fig.1D, E) suggesting that the localization of ESCRT-III/Vps4 to the PM might help to mitigate PM stress and prevent cell injuries.

Chronic inactivation of Ypk1 (chronic TORC2 inactivation is lethal) in *vps4*Δ mutants dramatically increased the frequency of PM integrity defects. This was assessed using propidium iodide (PI) staining. PI is membrane impermeable and only enters yeast cells when the integrity of the PM is corrupted. In WT cells, but also in *vps4*Δ and *ypk1*Δ single mutants the frequency of PI positive cells was low (Fig. 2E). However, the frequency of PI positive cells strongly increased in *vps4*Δ *ypk1*Δ double mutants (Fig. 2E), suggesting that they failed to maintain PM integrity.

These results demonstrated that TORC2-Ypk1 signaling and the ESCRT machinery helped to maintain the integrity of the PM. In response to acute PM stress the formation of ESCRT-III/Vps4 assemblies at the PM promoted cell survival. Chronic loss of Ypk1 signaling and the inability to form ESCRT-III/Vps4 assemblies at the PM compromised membrane integrity. This might be the leading cause for their synthetic genetic interaction and the mutual dependence of ESCRT function and TORC2-Ypk1 signaling

### TORC2-Ypk1 signaling prevents detrimental SL imbalance in ESCRT mutants

Next, we addressed how TORC2-Ypk1 signaling and the function of the ESCRT machinery converge to mitigate membrane stress. Therefore, we first defined the molecular nature of membrane stress in ESCRT mutants that is controlled by TORC2-Ypk1 signaling. Ypk1 is activated by TORC2 through phosphorylation primarily of Ser 644 and Thr 662 (Fig. 1C) (23, 69). Importantly, ESCRT mutants not only had slightly higher steady state Ypk1 protein levels, but also higher levels of active Ypk1 phosphorylated on Thr 662 (pT662) (Fig. 3A, Fig. S3A, B). We found no evidence for ESCRT-dependent lysosomal turnover of Ypk1 (70) in growing cells (Fig. S3C, D). The Ypk1 activation level in ESCRT mutants was substantial, but still lower compared to the hyperactivation of TORC2 signaling upon treatment with the SPT inhibitor myriocin (Fig. 3A, lane 1). Of note myriocin treatment for 150 min also reproducibly resulted in the up-regulation of Ypk1 protein levels (Fig. 3A, lane 1). The elevated Ypk1 protein levels that were detected in ESCRT mutants and in response to myriocin treatment might thus reflect Ypk1 stabilization by TORC2 phosphorylation as described earlier (69). We also observed a slight increase of *YPK1* mRNA in *vps4*Δ cells (Fig. S3E).

**Figure 3:**
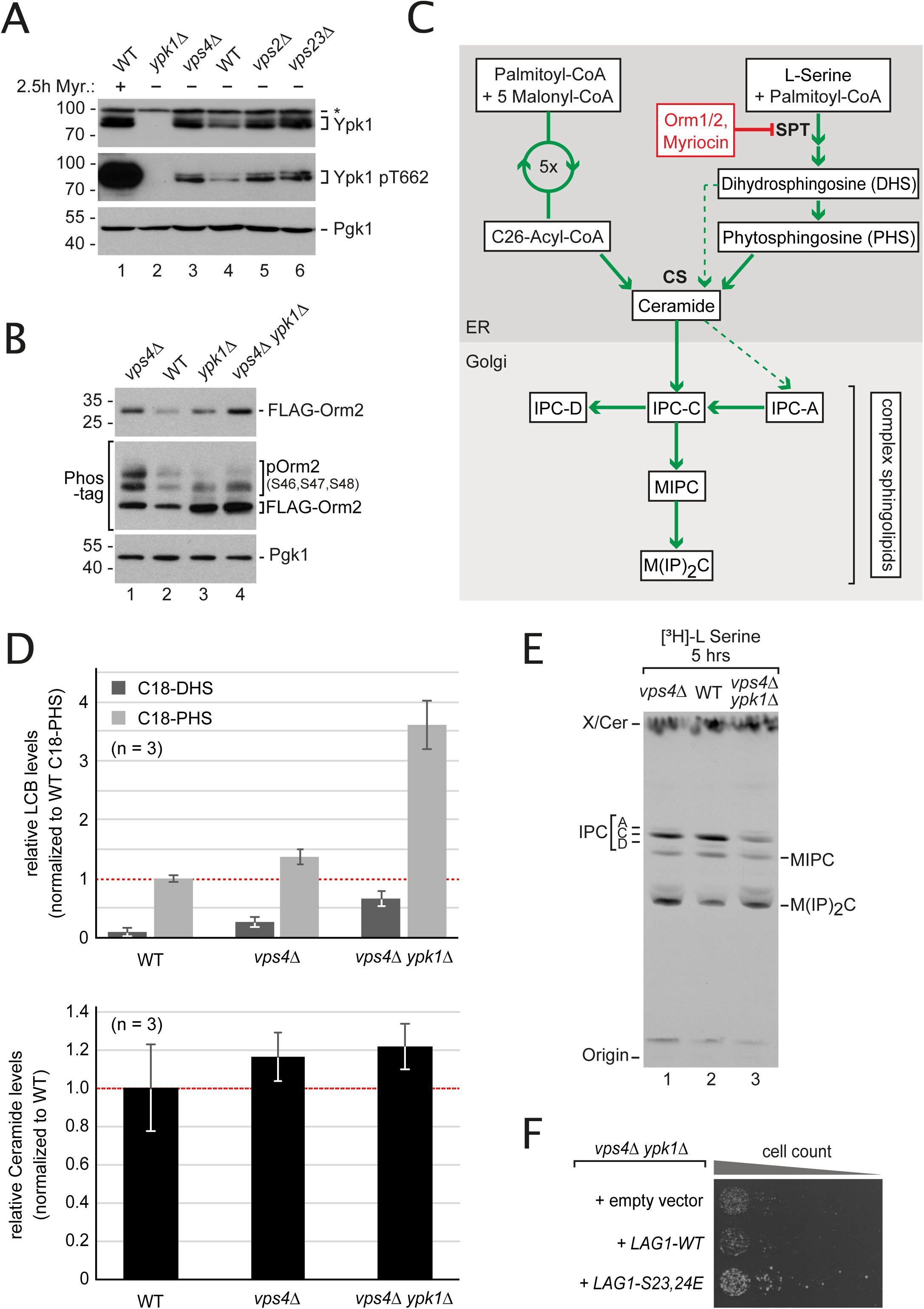
TORC2-Ypk1 signaling maintains sphingolipid homeostasis and prevents accumulation of long chain bases in ESCRT mutant cells. **(A)** SDS-PAGE and Western blot analysis with the indicated antibodies of total yeast lysates from WT cells and the indicated mutants. Myr., myriocin (1.5µM; 2.5 h) **(B)** SDS-PAGE or Phos-tag SDS PAGE and Western blot analysis with the indicated antibodies of total yeast lysates from WT cells and the indicated mutants expressing *FLAG-ORM2* (in *orm2*Δ background). **(C)** Schematic representation of SL biosynthesis in *S. cerevisiae*. SPT, serine-palmitoyl-coenzyme A transferase; CS, ceramide synthase; IPC, Inositolphosphorylceramide; MIPC, mannosylinositolphosphorylceramide; MIP_2_C, mannosyldiinositolphosphorylceramide. **(D)** The levels of the long-chain bases (LCB) C18-dihydrosphingosine (C18-DHS) and C18-phytosphingosine (C18-PHS) and of ceramides (presented as the sum of all species) in lipid extracts from WT cells, *vps4*Δ and *vps4*Δ *ypk1*Δ mutants were measured using LC-MS and quantified using non-yeast LCBs/ceramides as external standards. Data are normalized to WT levels of C18-PHS and ceramide (set to 1) and presented as mean ± standard deviation from 3 independent experiments. **(E)** Autoradiogram of sphingolipid extracts from [^3^H]-serine radiolabeled WT cells and the indicated mutants that were separated by thin layer chromatography. X indicates an unknown lipid species that is insensitive to myriocin treatment (data not shown). **(F)** Equal amounts of *vps4*Δ *ypk1*Δ cells expressing the indicated plasmids were spotted in serial dilutions on a YPD agar plate and incubated at 26°C. See also Figure S3.

Ypk1 has a range of different cellular targets (31,32,59). Among those, the paralogous ER proteins Orm1 and Orm2 are the essential targets of TORC2-Ypk1 (22, 31). In ESCRT mutant cells Orm2 protein levels and its Ypk1-dependent phosphorylation were increased (Fig. 3B). Deletion of *YPK1* reduced but did not completely abrogate steady state phosphorylation of Orm2 (presumably due to the presence of Ypk2), and caused upregulation of Orm2 protein levels (Fig. 3B). In *vps4*Δ *ypk1*Δ double mutants Orm2 protein levels were higher than in either single mutant.

TORC2-Ypk1 dependent phosphorylation of Orm2 and its paralogue Orm1 is required to de-repress the synthesis of LCBs by the SPT complex (15, 22) (Fig. 3C). TORC2 and Ypk1 also stimulate a subsequent step in the SL biosynthetic pathway by phosphorylating Lag1 and Lac1, two components of the ER-resident ceramide synthase (CS) (71, 72) (Fig. 3C). This concerted regulation avoids the toxic accumulation of biosynthetic intermediates and helps to maintain SL homeostasis (32).

We assessed the steady state levels of LCBs and ceramides using liquid chromatography - mass spectrometry (LC-MS) analysis. Despite the accumulation of Orm2 in *vps4*Δ cells, total LCBs (C18-PHS and C18-DHS) and ceramides were not lower than in WT cells (Fig. 3D). Also *in vivo* labeling of the SL pool with [^3^H]-L-serine in WT cells and *vps4*Δ cells showed only subtle differences in the steady state levels of mannosylated SL species, but the overall formation of [^3^H]-labelled complex SL was comparable (Fig. 3E). It seemed that steady state levels of SL synthesis in ESCRT mutants were maintained by Ypk1-dependent phosphorylation of Orm2. Consistently, loss of Ypk1 in *vps4*Δ mutants skewed SL biosynthesis. In *vps4*Δ *ypk1*Δ double mutants, the steady state LCB levels were increased (> 3-fold), while ceramide levels remained similar to *vps4*Δ single mutants (Fig. 3D). The loss of Ypk1 signaling in ESCRT mutants apparently caused an imbalance of SPT and CS activity, implying that the activity of CS might have become a bottleneck. Consistently, we observed that *vps4*Δ *ypk1*Δ double mutants produced less complex SL (predominantly inositolphosphoryl-ceramide (IPC)) than both WT and *vps4*Δ cells (Fig. 3E). Expression of a phospho-mimetic variant of CS can promote the formation of ceramides and complex SL independently of Ypk1 activity (32). Indeed, ectopic expression of the *LAG1*-S23,24E allele, which mimics constitutive Ypk1 phosphorylation (32), partially restored the growth of *vps4*Δ *ypk1*Δ cells (Fig. 3F).

These results demonstrate that ESCRT mutants have elevated Orm2 protein levels. TORC2-Ypk1 signaling partially compensates for the defects ensuing from Orm2 accumulation. Therefore, the SL profile of ESCRT mutants shows rather subtle differences compared to WT cells. Consistent with this notion, the deletion of *YPK1* in ESCRT mutants caused a detrimental imbalance in SPT and CS activity, characterized by an accumulation of LCBs and an impaired conversion into ceramides and complex SL. This imbalance in SL homeostasis might contribute to the defects in PM integrity observed in *vps4*Δ *ypk1*Δ mutants.

### In ESCRT mutants Orm2 is retained in the ER

While Orm2 protein levels were up-regulated in ESCRT mutants, the protein levels of Orm1 or other subunits of the SPOTS complex were unaffected (Fig. S4A). The changes in *ORM2* mRNA levels were minor and similar to *ORM1* (Fig. S4B), and cannot explain the increase of Orm2 protein. We concluded that the increase in Orm2 protein levels was caused by selective post-transcriptional mechanisms. The protein levels of Orm2 (but not Orm1) are controlled by a multistep process that delivers Orm2 from the ER to EGAD. First Orm2 is phosphorylated on Ser 46,47,48 by Ypk1. Next, phosphorylated Orm2 is exported from the ER in a COP-II-dependent manner. Once Orm2 arrives at the Golgi and on endosomes, it is recognized and ubiquitinated by the Dsc ubiquitin ligase complex, which ensures its proteasomal degradation (9).

To understand which of these steps are affected in ESCRT mutants, we compared them to EGAD mutants. Orm2 protein levels and its phosphorylation were similar in ESCRT (*vps4*Δ) and EGAD (*tul1*Δ) mutants (Fig. 4A). However, Orm2 accumulated in different subcellular compartments. In *tul1*Δ mutants GFP-Orm2 accumulated at the ER and on post-ER compartments, including Golgi, endosomes and the vacuolar limiting membrane, as reported earlier (9). In contrast, in the majority of *vps4*Δ cells, GFP-Orm2 was detected almost exclusively at the ER (Fig. 4B). Only in some cells, we observed a partial colocalization of GFP-Orm2 with the ESCRT substrate mCherry-CPS in class E compartments (Fig. S4C). Thus, it seemed that in ESCRT mutants Orm2 accumulated mainly at the ER.

**Figure 4:**
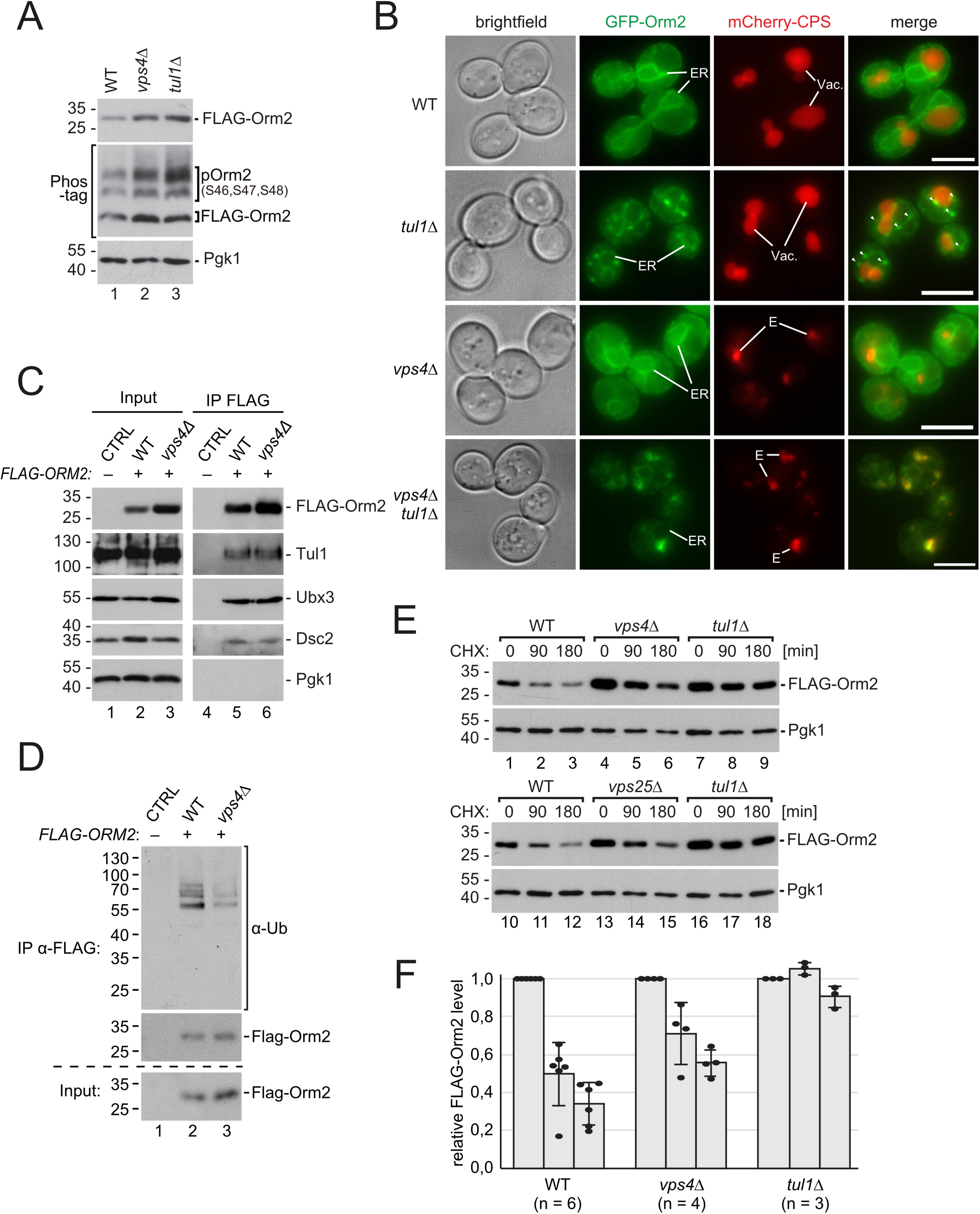
ER export of Orm2 is impaired in ESCRT mutants. **(A)** SDS-PAGE and Phos-tag SDS PAGE and Western blot analysis with the indicated antibodies of total yeast lysates from WT cells and the indicated mutants expressing *FLAG-ORM2*. **(B)** Epifluorescence and phase contrast microscopy of living WT cells and the indicated mutants expressing GFP-Orm2 WT (green) and mCherry-CPS (red). ER, endoplasmic reticulum; Vac, vacuole; E, class E compartment. Arrowheads indicate GFP-Orm2 accumulations in post-ER compartments. Scale bars 5µm. **(C)** SDS-PAGE and Western blot analysis with the indicated antibodies of input and elution (with FLAG peptide) from native FLAG-Orm2 immunoprecipitations (IP) from WT cells and *vps4*Δ mutants. Control cells expressed untagged Orm2. **(D)** SDS-PAGE and Western blot analysis with the indicated antibodies of input and elution (with FLAG peptide) from denaturing FLAG-Orm2 immunoprecipitations from WT cells and *vps4*Δ mutants. **(E)** WT cells and the indicated mutants were left untreated (0 min) or were treated with cycloheximide (CHX) to block protein synthesis for 90 min and 180 min at 26°C. Total cell lysates were analyzed by SDS-PAGE and Western blotting with the indicated antibodies. **(F)** Graphs display the FLAG-Orm2 protein levels determined by densitometric quantification of Western blots from cell lysates of WT cells and the indicated mutants 0, 90 and 180 min after the addition of CHX. Each experiment was repeated at least three times, FLAG-Orm2 levels were normalized to Pgk1 loading controls, and each time-point was related to t=0 min (set to 1). Data are presented as mean ±standard deviation. **(A-F)** All strains are in *orm2*Δ background. See also Figure S4.

Consistent with earlier reports (73), we detected Tul1 and Ubx3, two essential components of the Dsc complex, at the class E compartment in ESCRT mutants (Fig. S4D). Hence, the Dsc complex could degrade Orm2 once it reached the class E compartment. Indeed, GFP-Orm2 strongly accumulated in class E compartments when the EGAD pathway was additionally compromised in *vps4*Δ *tul1*Δ cells (Fig. 4B), in agreement with our earlier findings (9). The mutant Orm2-K25,33R, which is no longer ubiquitinated by the Dsc complex and hence no longer degraded, was also found predominantly in class E compartments in ESCRT mutants (Fig. S4E). Native immunoprecipitation (IP) of Flag-Orm2 demonstrated that a fraction of Orm2 still physically interacted with the Dsc complex in *vps4*Δ mutants (Fig. 4C), and denaturing IP of Flag-Orm2 showed that it was ubiquitinated (Fig. 4D), although to a lower extent compared to WT cells.

Consistently, Orm2 was still degraded in *vps4*Δ and *vps25*Δ mutants in cycloheximide chases, although the half-life was substantially longer (Fig. 4E). In WT cells Orm2 was degraded with a half-life of approximately 90 minutes, and in *vps4*Δ mutants the half-life of Orm2 was approximately 180 minutes (Fig. 4E, F). In contrast, Orm2 degradation was fully blocked in mutants of the Dsc complex (*tul1*Δ) (9) (Fig. 4E, F).

Collectively, these results support the idea that the ER export of Orm2 is reduced in ESCRT mutants and causes accumulation of Orm2, despite a functional EGAD machinery.

### ER accumulation of Orm2 is caused by calcineurin and renders ESCRT mutants addicted to TORC2-Ypk1 signaling

Our results so far present a conundrum: In ESCRT mutants Orm2 is phosphorylated by TORC2-Ypk1 (Fig. 3B, Fig. 4A), but still accumulates at the ER. Mutation of Ser 46,47,48 to alanine in Orm2 (Orm2-3A) prevented phosphorylation by Ypk1 (22), reduced Orm2 ER export and subsequent EGAD (9), and therefore Orm2-3A accumulated in WT cells and in *vps4*Δ mutants (Fig. 5A). We also reported earlier that Ser 46,47,48 to aspartic acid mutations (Orm2-3D) mimic constitutive phosphorylation and trigger constitutive ER export, leading to degradation via EGAD. Therefore, Orm2-3D protein levels are low in WT cells. Importantly also in *vps4*Δ mutants, Orm2-3D protein levels were strongly reduced (Fig. 5A). This result implied that in *vps4*Δ mutants the phospho-mimetic Orm2-3D mutant could be exported more efficiently from the ER, and then was degraded by EGAD. Remarkably, expression of the Orm2-3D mutant, but not Orm2-3A, improved also the growth of *vps4*Δ *ypk1*Δ cells (Fig. 5B).

**Figure 5:**
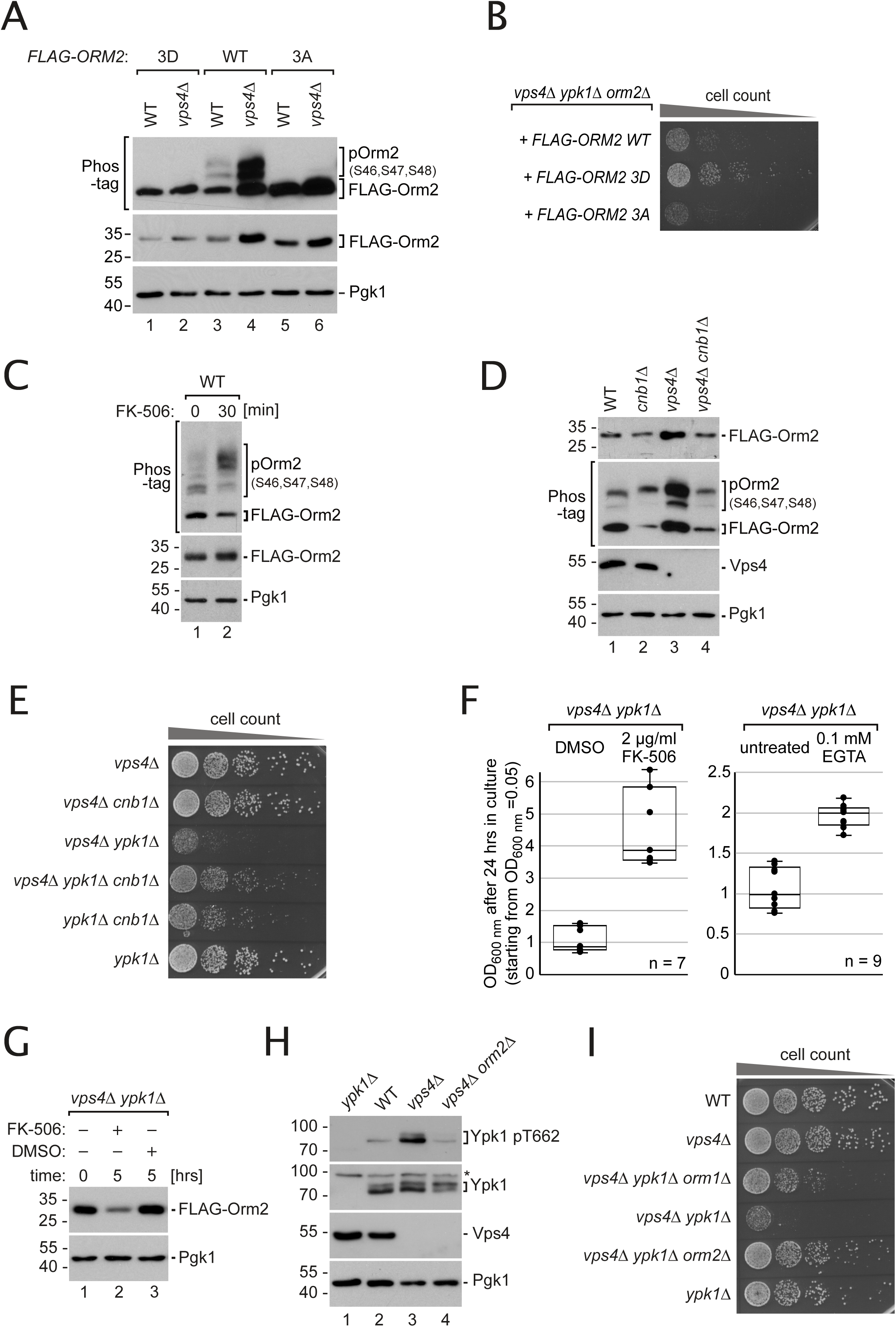
Orm2 accumulation causes membrane stress in ESCRT mutants. **(A)** SDS-PAGE or Phos-tag SDS PAGE and Western blot analysis with the indicated antibodies of total yeast lysates from WT or *vps4*Δ cells expressing *FLAG-ORM2-WT, -3A or -3D* (in *orm2*Δ background). **(B)** Equal amounts of *vps4*Δ *ypk1*Δ *orm2*Δ cells expressing the indicated *FLAG-ORM2* plasmids were spotted in serial dilutions on a auxotrophic selection medium agar plate and incubated at 26°C. **(C)** SDS-PAGE and Phos-tag SDS PAGE and Western blot analysis with the indicated antibodies of total yeast lysates from WT cells expressing *FLAG-ORM2* (in *orm2*Δ background) treated with FK-506 (2 µg/ml) for the indicated time. **(D)** SDS-PAGE and Phos-tag SDS PAGE and Western blot analysis with the indicated antibodies of total yeast lysates from WT cells and the indicated mutants expressing *FLAG-ORM2*. **(E)** Equal amounts of the indicated single, double or triple mutants in serial dilutions were incubated on a YPD agar plate at 26°C. **(F)** Left: Growth of *vps4*Δ *ypk1*Δ cells in YPD medium in presence of 2 µg/ml FK-506 or vehicle (DMSO). Right: Growth of *vps4*Δ *ypk1*Δ cells in YPD medium in presence of 0.1 mM EGTA or untreated. Cells were inoculated to OD_600nm_ = 0.05 and grown for 24 hours in 3 independent experiments. The circles indicate the individual measurements (7 and 9 technical replicates). **(G)** SDS-PAGE PAGE and Western blot analysis with the indicated antibodies of total yeast lysates from *vps4*Δ *ypk1*Δ cells expressing *FLAG-ORM2* (in *orm2*Δ background) treated with FK-506 (2 µg/ml) or vehicle (DMSO) for the indicated time. **(H)** SDS-PAGE and Western blot analysis with the indicated antibodies of total yeast lysates from WT cells and the indicated mutants. **(I)** Equal amounts of WT cells and indicated single, double or triple mutants in serial dilutions were incubated on a YPD agar plate at 26°C. See also Figure S5.

Since mimicking of constitutive phosphorylation improved Orm2 ER export and degradation in ESCRT mutants, we can rule out general defects in ER export machineries or Orm2 degradation. Rather, these results implied that untimely dephosphorylation might hamper ER export of Orm2. The calcium-dependent protein phosphatase calcineurin is a major antagonist of TORC2 signaling (74), with regulatory roles in sphingolipid biosynthesis (26,32,60). Moreover, membrane stress during heat shock has been linked to a mis-regulation of calcium dynamics and calcineurin signaling impairing sphingolipid synthesis in the ER (60). ESCRT mutant cells have been reported to be prone to accumulate intra-cellular calcium (75), with increased activity of a calcineurin-dependent transcriptional reporter (76). Therefore, we hypothesized that calcineurin activity in ESCRT mutants could counteract TORC2-Ypk1 signaling and thereby delay ER export and degradation of Orm2.

To test this hypothesis, we first acutely inhibited calcineurin with FK-506. Inhibition of calcineurin for 30 minutes resulted in hyper-phosphorylation of Orm2 (Fig. 5C). It had been shown before that calcineurin inhibition does not augment Ypk1 activation by TORC2 or Pkh1/2 (22,24,32). Hyper-phosphorylation of Orm2 in response to calcineurin inhibition suggested that this phosphatase was involved in the dephosphorylation of Orm2. Next, we deleted the regulatory calcineurin subunit Cnb1, which chronically abrogates calcineurin activity (77). Chronic loss of calcineurin activity in *cnb1*Δ cells also caused hyper-phosphorylation of Orm2 and additionally led to slightly lower Orm2 protein levels compared to WT cells. In *vps4*Δ mutants, deletion of *CNB1* markedly decreased Orm2 to levels that were similar to WT cells (Fig. 5D). Thus, calcineurin activity results either directly or indirectly in the dephosphorylation of Orm2, and thereby antagonizes TORC2-Ypk1-induced ER export and degradation of Orm2.

Calcineurin activity also rendered ESCRT mutants dependent on TORC2-Ypk1 signaling, since deletion of *CNB1* partially restored the growth of *vps4*Δ *ypk1*Δ mutants (Fig. 5E). Likewise, pharmacological inhibition of calcineurin activity (2µg/ml FK-506) or addition of the calcium-chelator EGTA to the growth medium improved growth of *vps4*Δ *ypk1*Δ cells (Fig. 5F). Calcineurin inhibition with FK-506 markedly decreased Orm2 protein levels in the *vps4*Δ *ypk1*Δ double mutants (Fig. 5G). The inhibitory function of calcineurin was likely independent of the transcription factor Crz1 which is a major downstream effector of calcineurin (78, 79). Unlike the deletion of *CNB1*, deletion of *CRZ1* did not rescue growth of *vps4*Δ *ypk1*Δ cells (Fig. S5A). This is consistent with a role of calcineurin in controlling either directly or indirectly Orm2 phosphorylation. Thus, calcineurin activity in ESCRT mutants reduced the ER export and degradation of Orm2, which contributed to accumulation of Orm2 in the ER and rendered the growth of ESCRT mutants dependent on TORC2-Ypk1 signaling.

Clearly, the accumulation of Orm2 constituted a major membrane stress factor in ESCRT mutants: In *vps4Δ orm2*Δ double mutants Ypk1 activation by TORC2 (pT662) was no longer elevated (Fig. 5G). In addition, *vps4Δ orm2*Δ cells grew better in presence of PalmC than *vps4*Δ single mutants, suggesting a decreased susceptibility to PM stress (Fig. S5B). Furthermore, deletion of *ORM2* in *vps4*Δ *ypk1*Δ double mutants restored membrane integrity and decreased the fraction of PI-positive cells to *orm2*Δ single mutant levels (Fig S5C). Finally, the deletion of *ORM2* fully rescued the growth defect of *vps4*Δ *ypk1*Δ cells (Fig. 5H). A partial rescue was also observed upon deletion of *ORM1*.

Our results suggest that calcineurin activity in ESCRT mutant cells hampered the homeostatic regulation of Orm2 ER export. In turn, Orm2 accumulated at the ER and was no longer efficiently degraded by EGAD. The accumulation of Orm2 at the ER renders the growth of ESCRT mutants dependent on TORC2-Ypk1 signaling to prevent imbalance in SL homeostasis and loss of plasma membrane integrity (Fig. S5D).

## Discussion

Here we show that the ESCRT machinery contributes directly and indirectly to PM homeostasis in budding yeast and thereby takes part in ER-PM crosstalk to preserve the barrier function of the PM.

ESCRT-III/Vps4 assemblies that form at the plasma membrane might play a direct role in maintaining membrane integrity. A role for ESCRT complexes in repairing damaged PM was described in human cells (56,66–68). There, injury-triggered calcium increase results in assembly of ESCRT-III and accessory proteins at the site of repair via Alix (68). Yet, it is unclear if ESCRTs also prevents membrane lesions in the first place. We observed ESCRT assemblies at the PM in response to indirect (i.e by reduction of membrane tension) and direct TORC2 inhibition. The major ESCRT-III subunit Snf7 and the AAA-ATPase Vps4 were detected at PM invaginations and subcortical foci, which are reminiscent of stalled endocytic buds. This seems plausible, since TORC2 promotes endocytosis by controlling actin and endocytic proteins and by modulating membrane tension (31,36,65).

How ESCRT-III was recruited to these structures is currently unclear, but several scenarios appear possible. ESCRT-III/Vps4 recruitment could be driven by ESCRT-0, -I or -II complexes interacting with the ubiquitinated membrane proteins in these stalling endocytic buds. Indeed, ESCRT-II mutants also genetically interact with *YPK1* (Fig. S1B). Alternatively, Bro1 – like Alix in human cells - might help to recruit ESCRT-III directly to the PM. In budding yeast, ESCRT-III subunits (Snf7) are also recruited to the PM during adaptation to alkaline pH and are critically required for the regulator of Ime2 (RIM) pathway (80). Alternatively, changes in PM tension might recruit the ESCRT machinery as described (81, 82). Interestingly, ESCRT-III subunits are recruited onto endosomal membranes when membrane tension is reduced (82). In analogy, we speculate that ESCRT assemblies might recognize areas of decreased tension at the PM.

Once there, the ESCRT machinery is involved in consolidating a slack PM. This might be achieved either through stabilization of PM deformations, or via active remodeling (e.g. removing (budding out) or deforming membranes), which could increase tension again. How exactly ESCRT-III/Vps4 assemblies help to maintain the integrity of the PM in response to tensile stress remains an open question at the moment.

The results from our genetic screen indicated that ESCRT-deficient cells are particularly sensitive to perturbations in the homeostasis of two classes of lipids: the fungal cholesterol analog ergosterol and sphingolipids. In concert, these lipids are known to promote membrane rigidity and also to stabilize many membrane proteins (83–86). In yeast, they are important for the formation of eisosomes (87), protein-coated membrane invaginations that have been suggested to function as membrane reservoirs that can balance changes in membrane tension, in analogy to caveolae (88–90). Moreover, eisosomes are involved in the regulation of TORC2 signaling (24). Sphingolipids and ergosterol might therefore become essential in ESCRT mutants to counteract membrane stress.

Orm2 emerges here as a central player in membrane homeostasis. The accumulation of Orm2 at the ER in ESCRT mutants and/or in post-ER compartments in EGAD mutants causes membrane stress in budding yeast, judged from increased activity of TORC2 (9). Hence maintaining proper control of Orm2 levels is important to restrain SPT activity and prevent the potentially toxic accumulation of metabolic intermediates, such as LCBs (91). Similarly, failure to control ORMDL protein levels is associated with several disease conditions in humans (15,18–21), which are probably connected to mis-regulation of SPT activity (92, 93) and might well involve increased susceptibility to membrane stress.

Levels of LCBs are usually tightly controlled on the levels of synthesis, degradation and conversion into ceramides and complex SL (94, 95). Primarily Ypk1 (and to a lesser extent Ypk2) regulates flux of LCBs into complex SL through phosphorylation of the CS subunits Lag1 and Lac1 (32). Failure to phosphorylate CS also causes accumulation of LCBs and diminished formation of complex SL. Thus, we speculate that in *vps4*Δ *ypk1*Δ cells the coordination of SPT and CS activity is lost. Probably SPT activity can still be to some degree de-repressed by TORC2 via Ypk2, whereas CS activity cannot be efficiently stimulated when Ypk1 is deleted (32). This may additionally be aggravated by the accumulation of Orm2 in the ER, because high Orm2 levels can diverge Ypk1 (and hence probably also Ypk2) activity away from other important targets (32). Collectively, this leads to a built up of LCBs in *vps4*Δ *ypk1*Δ cells and the ensuing defects in membrane integrity. Consistently, mimicking phosphorylation of Lag1 slightly improved the growth of *vps4*Δ *ypk1*Δ mutants (Fig. 3F).

The major culprit for the accumulation of Orm2 at the ER in ESCRT mutants appears to be its premature dephosphorylation in a calcineurin-dependent manner, which decreases ER export of Orm2 and thus slows down Orm2 degradation through the EGAD pathway. Calcineurin is activated by elevated intracellular calcium levels, and calcineurin activity is probably increased in ESCRT mutants (76). ESCRT mutants have been found to be sensitive to calcium, which can be suppressed by calcineurin inhibition or by stimulating intracellular calcium storage (75, 96). *vps4*Δ mutants upregulate the Golgi calcium pump Pmr1 (97), possibly to decrease intracellular calcium. Pmr1 is also essential for the survival of ESCRT mutants (9). Similarly, Csg2 is required for growth under high calcium conditions (98) and for growth of *vps4*Δ mutants (Fig. S1A), indicating tight links between the ESCRT machinery, calcium signaling and the regulation of SL biosynthesis.

The growth defect of *vps4*Δ *ypk1*Δ double mutants was alleviated by calcineurin inhibition and extracellular calcium chelation, suggesting that increased calcium uptake drives detrimental calcineurin activity in these cells. Increased calcium influx in ESCRT mutants may be caused by deregulated trafficking or turnover of the plasma membrane calcium channel (Cch1/Mid1), which is likely a substrate of the ESCRT machinery. Mid1 could also respond to increased membrane stress in ESCRT mutants, since Mid1 is a stretch-activated channel and allows calcium influx and calcineurin signaling to respond to mechanical membrane stress (99). Alternatively, ER-PM contact sites may be corrupted in ESCRT mutants. The disruption of these contact sites leads to increased calcium influx and calcineurin activity, which increased LCBs and decreased ceramide levels (60).

At the moment, it is unclear whether calcineurin directly dephosphorylates Orm2. The sequence of Orm2 does not contain an obvious calcineurin binding site (100, 101). The Cdc55-containing protein phosphatase 2A (PP2A) controls Orm2 phosphorylation levels during heat shock (102). Inactivation of Cdc55-PP2A or another phosphatase upon calcineurin-inhibition cannot be ruled out as an indirect cause leading to dephosphorylation of Orm2, but to our knowledge has not been reported. However, since calcineurin antagonizes TORC2 and Ypk1 activity also towards other targets (29,32,74,103), it might also directly contribute to the phosphorylation status of Orm2. In addition to lowering Orm2 protein levels, inhibition of calcineurin is also expected to promote phosphorylation of the CS subunits Lag1/Lac1 and thereby formation of ceramides, which mitigates the buildup of harmful LCBs (32). Thus, calcineurin inactivation probably has several beneficial effects in ESCRT mutants, which collectively render their growth largely independent of Ypk1 signaling.

In summary, we propose that ESCRT function maintains PM homeostasis on several levels: (i) through the MVB pathway by degrading membrane proteins and preventing proteotoxic stress (104), (ii) by preserving or repairing the PM in response to membrane stress, and (iii) by maintaining calcium homeostasis, which is important for controlled output of TORC2-Ypk1 signaling and regulation of Orm2. The latter promotes biosynthetic SL flux and thereby preserves PM homeostasis. In ESCRT mutants, regulation of Orm2 is disturbed because of increased calcineurin-dependent dephosphorylation, and hence its ER export and turnover through EGAD is impaired. The buildup of Orm2 protein levels in the ER constitutes a stress that must be counteracted through increased TORC2-Ypk1 activity (Fig. S5D). As long as this is possible, ESCRT mutants have little fitness defects. However, if phosphorylation of Orm2 is additionally impaired by any means (deletion of *YPK1*; inhibition of TORC2), or if plasma membrane homeostasis is acutely challenged, ESCRT mutants are no longer able to compensate. Now their PM integrity becomes compromised, and viability decreases dramatically. Importantly, the mutual dependence of the ESCRT machinery and TORC2-Ypk1 signaling was broken by reducing steady state levels of Orm2, either through deletion of the gene, or by enforcing phosphorylation, ER export and degradation.

These findings demonstrate the importance of post-translational regulation of ORMDL family proteins for membrane homeostasis, and establish ORMDL protein levels as a central determinant of PM stress in eukaryotic cells. Our data connect the ESCRT machinery to PM stress and deregulated calcineurin signaling, and explain how ESCRT mutants become addicted to TORC2-dependent signaling.

## Experimental Procedures

### Yeast strains, plasmids and growth conditions

Genetic modifications were performed by PCR and/or homologous recombination using standard techniques. Plasmid-expressed genes including their native promoters and terminators were amplified from yeast genomic DNA and cloned into centromeric vectors (pRS series) (105). Tagged version of Orm1 and Orm2 (3xHA, 3xFLAG, or GFP) and their respective mutants were expressed from plasmids in the respective deletion mutants replacing the endogenous protein (with exception of Fig. 5D, where *FLAG-Orm2* was co-expressed). All constructs were analyzed by DNA-sequencing and transformed into yeast cells using standard techniques. Genotypes of yeast strains and plasmids used in this study as well as primer for PCR-based genetic modifications and cloning are listed in Table S2.

All *S. cerevisiae* strains in this study were SEY6210 derivatives. For liquid cultures, cells were incubated in YNB synthetic medium supplemented with amino acids (according to respective auxotrophies) and 2% glucose at 26°C in a shaker and were grown to midlog phase (OD_600_=0.5 – 0.8). For growth on agar plates, yeast cells were diluted to OD_600nm_ = 0.05 and spotted in serial dilutions on YPD or YNB (auxotrophic selection medium) plates at the indicated conditions. Rapamycin (Sigma) was added at the indicated concentrations from a 1 mM stock in DMSO and myriocin (Sigma) was added to 1.5 µM from a 5 mM stock in methanol. Untreated controls were supplied with the appropriate amount of the respective solvent. For the assessment of growth in presence of PalmC, cells were grown into log phase, diluted to an OD_600nm_ of 0.1 into fresh YNB medium containing PalmC (Sigma) at the indicated concentration (from a 5 mM stock in DMSO) and grown at 26°C, 180 rpm. For the assessment of growth in presence of FK-506 (Sigma; stock 20 mg/ml in DMSO) or EGTA (Sigma; stock 500 mM in water) cells were grown into log phase, diluted to an OD_600nm_ of 0.05 into fresh YPD medium containing 2 µg/ml FK-506 or 0.1 mM EGTA. After 24 hours the OD_600nm_ was measured.

### Analysis of genetic interaction data

Gene Ontology (GO) enrichment analysis (106) was performed using the 119 genes listed in Appendix Table S1 of (9) as synthetically lethal with *vps4*Δ in two biological replicates or lethal in one and sick in the other replicate. They were analyzed with the ‘generic GO-Slim: process’ terms using the GO Slim mapper of the *Saccharomyces* genome database. GO term fusion and enrichment analysis was performed as described (9). Only significantly scored GO terms (p < 0.05) are presented in Fig. 1A. The full analysis is presented in Table S1.

### Preparation of whole cell protein extracts, Western blot analysis and immunodetection

To prepare whole cell lysates, proteins were extracted by alkaline extraction (107). When protein phosphorylation was also analyzed, extraction was done by a modified protocol with phosphatase inhibition as described (9). Protein extracts were denatured in Lämmli sample buffer, separated by SDS-PAGE (Biorad Mini Protean) and transferred to PVDF membranes by semi-dry electro-blotting. Phos-tag SDS PAGE as well as immunodetection of ubiquitinated Orm2 were done as described (9). Antibodies used in this study are listed in Table S2.

### FLAG-Orm2 immunoprecipitations

Immunoprecipitations of FLAG-Orm2 under denaturing or non-denaturing lysis conditions were done as described in detail in (9).

### RNA isolation and quantitative PCR (RT-qPCR)

RNA isolation, cDNA synthesis and quantitative RT-PCR analysis were performed as described previously (9, 108). TAQman gene expression assays were from Thermo Fisher (*YPK1*: Sc04141261_s1; *ORM1*: Sc04125000_s1; *ORM2*: Sc04149509_s1; housekeeping gene *PGK1*: Sc04104844_s1). RT-qPCR analysis was done from 4-6 independent biological samples and each in 3-4 technical replicates. Data were analyzed with the PikoReal software (version 2.2; Thermo Scientific) with manual threshold adjustment, and relative mRNA abundance was calculated in Microsoft Excel (Version 16.16.2; RRID:SCR_016137) using the ΔΔC_T_ method (109).

### Fluorescence live cell wide field microscopy

For microscopy, cells were grown to midlog (OD_600_=0.5 – 0.8) phase in YNB media, concentrated by centrifugation and directly mounted onto glass slides. For imaging of the Dsc complex (Fig. S4B), cells were grown for 36 hours on selective YNB agar plates, dissolved in sterile water and mounted for imaging. Live cell wide field fluorescence microscopy was carried out using a Zeiss Axio Imager M1 equipped with a sola light engine LED light source (Lumencore), a 100x oil immersion objective (NA 1.45,) standard GFP and mCherry fluorescent filters, a SPOT Xplorer CCD camera, and Visitron VisiView software (version 2.1.4). The brightness and contrast of the images in the figures were adjusted using Photoshop CS5 (Adobe Version 12.0.4x64; RRID:SCR_014199). For merged images the levels of red and green channels were separately adjusted.

### Assessment of plasma membrane integrity by propidium iodide staining

Cells were grown in YPD medium and kept in logarithmic growth phase for >12 hours. 1 ml of culture was supplemented with 3 mg/ml propidium iodide (Sigma; stock 1 mg/ml in phosphate buffered saline (PBS)) and incubated at RT for 10 minutes. Cells were recovered by centrifugation (10000xg, 4°C, 2 min), washed once with 1 ml ice cold PBS, resuspended in 0.5 ml ice cold PBS and kept on ice. Propidium iodide-positive cells were counted on an Attune™ NxT Acoustic Focusing Cytometer (Life Technologies) with Attune™ NxT software (v. 3.1.1243.0). Debris and cell doublets were excluded. Propidium iodide signal was measured with 488 nm excitation and emission in the 695/40 nm window. In each experiment 50000 cells of each genotype were counted. Gating was adjusted so that more than 99.9% of WT cells were propidium iodide-negative.

### Immuno-gold electron microscopy

As previously described (44) samples were high-pressure frozen, freeze-substituted and rehydrated (110), followed by indirect immunogold labeling of 100 nm-thick, thawed cryosections (44). Antibodies used were goat polyclonal anti GFP (1:500; Rockland) and rabbit anti goat Fab’ NANOGOLD™ (1:150; Nanoprobes). Transmission electron microscopy was performed on a Philips CM120 (now: Thermo Fisher Scientific).

### Cycloheximide chase assay

Logarithmically growing cells (20 OD_600nm_) were harvested by centrifugation and cells were resuspended in fresh medium to concentration of 0.4 OD/ml. 10 ml (4 OD_600nm_) were immediately (t = 0 min) harvested by centrifugation, washed once with ice-cold 10 mM NaF solution, and pellets were snap frozen in liquid nitrogen. To the remaining culture 50 µg/ml cycloheximide (Sigma Aldrich) was added from a 10 mg/ml stock. After the indicated time points 10 ml culture were harvested, washed and frozen as above. Whole cell extracts were prepared by alkaline extraction. SDS-PAGE, Western blot detection and quantification were done as described (9).

### Metabolic sphingolipid labeling, sphingolipid extraction and thin layer chromatography

Sphingolipid labeling and extraction were done as described previously (9, 111). Logarithmically growing cells (5 OD_600nm_) were harvested by centrifugation and resuspended in 470 µl fresh medium. 30 µCi of [^3^H]-L-serine (1µCi/µl; Hartmann Analytic) was added and cells were incubated at 30°C, 700 rpm for 5 hours. Proteins were precipitated with 4.5% perchloric acid, cell pellets were washed with ice-cold 100mM EDTA, resuspended in 50µl water, and subjected to mild alkaline methanolysis (50% methanol; 10% 1-butanol; 10% monomethylamine; 50 minutes at 50°C) of ester lipids. Lysates were vacuum dried in a speed vac (37°C), pellets were resuspended thoroughly in 300 µl water by sonication, and tritium-incorporation of was assessed for normalization in duplicates by szintillation counting (LS6500, Beckmann Coulter). Sphingolipids were extracted 3 times with water-saturated 1-butanol, vacuum dried in a speed vac (37°C), and dissolved in 50 µl chloroform : methanol : water 10:10:3 for thin layer chromatography. Thin-layer chromatography (solvent system chloroform : methanol : 4.2N ammonium hydroxide 9:7:2) was done as described (111) on aluminium silica TLC plates (Sigma) for 75 minutes. Subsequently, plates were dried under air flow and treated twice with En^3^Hance autoradiography enhancer (Perkin-Elmer), dried again, and exposed to autoradiography films (CL-Xposure Film, Thermo) at −80°C. The identity of labeled bands was confirmed with specific mutant yeast strains or inhibitors (data not shown). A tritium-labeled band X close to the solvent front that was observed previously (111) is not sensitive to myriocin treatment (Sigma Aldrich, 1.5µM), suggesting that it is not a sphingolipid.

### Lipid extraction and liquid chromatography - mass spectrometry (LC-MS)

For combined LCB and ceramide analysis (Fig. 3D) lipids were extracted from lysed yeast cells according to 200 µg of protein by chloroform/methanol extraction (112). Prior to extraction an internal standard mix containing sphingosine (LCB 17:0) and ceramide (CER 18:0/17:1) was spiked into each sample for normalization. Quantification was done by external quantification with the same lipids in different concentrations. Dried lipid samples were dissolved in a 65:35 mixture of mobile phase A (60:40 water/acetonitrile, including 10 mM ammonium formate and 0.1% formic acid) and mobile phase B (88:10:2 2-propanol/acetonitrile/H_2_0, including 2 mM ammonium formate and 0.02% formic acid). HPLC analysis was performed employing a C30 reverse-phase column (Thermo Acclaim C30, 2.1 x 250 mm, 3 µm, operated at 40° C; Thermo Fisher Scientific) connected to an HP 1100 series HPLC (Agilent) HPLC system and a QExactive*PLUS* orbitrap mass spectrometer (Thermo Fisher Scientific) equipped with a heated electrospray ionization (HESI) probe. The elution was performed with a gradient of 20 min; from 0–1 min elution starts with 40% B and increases to 100% in a linear gradient over 13 min. 100% B is maintained for 3 mins. Afterwards solvent B was decreased to 40% and maintained for another 3.8 minutes for column re-equilibration. The mass spectrometer was run in negative and positive ion mode. The scan rate was set from 200-1200 m/z. Mass resolution was 70,000 with an AGC target of 3,000,000 and a maximum injection time of 100 ms. The MS was operated in data dependent mode. For MS/MS the resolution was 35000 with a maximum injection time of 50 ms and an AGC target of 100,000. The loop count was ten. Selected ions were fragmented by HCD (higher energy collision dissociation) with a normalized collision energy of 30. The dynamic exclusion list was set to 10s to avoid repetitive sequencing. Ceramide peaks were identified using the Lipid Search algorithm (MKI, Tokyo, Japan). Peaks were defined through raw files, product ion, and precursor ion accurate masses. Ceramides were identified by database (>1,000,000 entries) search of negative ion adducts. LCBs were identified by positive ion adducts. The accurate mass extracted ion chromatograms were integrated for each identified lipid precursor and peak areas obtained for quantitation. Internal standards were used for normalization and an external standard curve was used to calculate absolute values (in pmol/µg protein). Ceramides are presented as the sum of all quantified ceramide species. For comparison, data were normalized to WT levels (set to 1) of ceramide and C18-PHS, respectively. Data are presented as mean ±standard deviation from three independent experiments.

### Quantification and statistical analysis

Statistical details and sample numbers of quantitative analyses can be found in the respective figures and corresponding figure legends. Quantitative data is usually displayed as mean ±standard deviation from at least 3 biological replicates and relates to the wildtype control.

#### (i) GO analysis of genetic interaction data

A hypergeometric test was used to estimate if the mapped GO term is significantly enriched with the selected genes. The null hypothesis is that the selected genes are randomly sampled from all yeast genes. The resulting p-values were corrected with the Benjamini-Hochberg method. All adjusted p-values below 0.05 were reported.

#### (ii) Quantification of Snf7-eGFP and Vps4-eGFP localization

Images were taken in an unbiased manner by selecting and focusing cells only in brightfield view. 50 cells of each genotype and condition were imaged and visually inspected for the presence and number of eGFP puncta at the cell cortex. Cells were grouped into cohorts of 0-2, 3-5 and more than 5 cortical eGFP puncta.

#### (iii) Quantification of cell growth

Cell growth after 2 hours in culture was measured photometrically by absorbance at 600 nm. Measurements from 3 independent experiments (each with up to three parallel cultures representing technical replicates) were presented with box-whisker plots indicating the median, first and third quartile. Individual data points including outliers were also plotted.

#### (iv) Quantification of Western Blot analysis

Western blot signals were quantified by densitometry using ImageJ2 (Version 2.0.0-rc49/1.51h; RRID:SCR_003070) (113), quantifications were exported to Microsoft Excel (Version 16.16.2; RRID:SCR_016137), normalized to the respective Pgk1 loading controls, and presented as mean ±standard deviation from at least three independent experiments. t=0 was set to 1.

#### (v) Quantitative PCR (RT-qPCR)

Mean, normalized ΔΔC_T_ values were individually calculated for four independent biological replicates (each with 3-4 technical replicates), log2-transformed to calculate fold change, and presented as mean fold change over WT ±standard deviation. A one-sided Student’s t-test was used on the mean ΔΔC_T_ values (i.e. prior to log transformation) of the four biological replicates of to assess statistical significance.

#### (vi) Lipid extraction and mass spectrometry

Internal standards for ceramides (Cer 18:1;2/17:0;0) and LCBs (LCB 17:0) spiked in prior to extraction were used for normalization and an external standard curve was used to calculate absolute values (in pmol/µg protein). To compare data from different experimental setups, data were normalized to WT levels of the most abundant LCB and ceramide species (C18-phytosphingosine and Cer44:0;4). Data are presented as mean ±standard deviation from three independent experiments.

## Data availability

All relevant data has been included in the paper in main figures and supplemental information.

## Acknowledgments

We would like to thank Theresa Dunn, Peter Espenshade, Ming Li and Howard Riezman for providing reagents, and Jennifer Kahlhofer and Manuel Haschka for help with flow cytometry.

## Author contributions

O.S. performed the SL-TLC analyses and growth experiments; O.S., Y.W., S.S., M.A.W., V.B., R.L. and C.S. constructed yeast strains and plasmids and conducted microscopy, biochemical and genetic experiments; M.A. performed the statistical analysis of the GO dataset; M.H. performed electron microscopy; S.E. and F.F. analyzed ceramide and LCB levels; O.S., S.S. and D.T. analyzed data; O.S. and D.T. guided the story; O.S. and D.T. wrote the manuscript with input from all authors.

## Funding and additional information

Work in the Teis laboratory was supported by the Austrian Science Fund (FWF-Y444-B12, P30263, P29583) and MCBO (W1101-B18) to D.T and EMBO/Marie Curie (ALTF 642-2012; EMBOCOFUND2010, GA-2010-267146), MUI-START (2013042023) and ‘Tiroler Wissenschaftsfond’ to O.S. The Stefan laboratory is supported by MRC funding to the MRC LMCB University Unit at UCL, award code MC_UU_00012/6.

## Abbreviations and nomenclature

AAA: ATPase associated with diverse cellular activities
CS: ceramide synthase
DHS: dihydrosphingosine
Dsc: defective in SPRBP cleavage
EGAD: endosome and Golgi-associated degradation
EGTA: Ethylene glycol-bis(2-aminoethylether)-N,N,N′,N′-tetraacetic acid
ER: endoplasmic reticulum
ERAD: endoplasmic reticulum-associated degradation
ESCRT: endosomal sorting complex(es) required for transport
GO: gene ontology
HMGR: 3-hydroxy-3-methylglutaryl coenzyme A reductase
IP: immunoprecipitation
IPC: inositolphosphoryl ceramide
LCB: long chanin base
LC-MS: liquid chromatography – mass spectrometry
MVB: multivesicular body
Orm2-3A: Orm2 serine 46,47,48 to alanine mutant (non-phosphorylatable)
Orm2-3D: Orm2 serine 46,47,48 to aspartic acid mutant (phospho-mimetic)
ORMDL: Orosomucoid-like protein (mammalian Orm1/2 paralogues)
SGK1: serum and glucocorticoid-induced kinase 1
PalmC: palmitoyl carnitine
PHS: phytosphingosine
PI: propidium iodide
PM: plasma membrane
RIM: regulator of Ime2
SL: sphingolipid
SPOTS: SPT-Orm1/2-Tsc3-Sac1 complex’
SPT: serine-palmitoyl-coenzyme A transferase
SREBP: sterol-responsive element binding protein
TORC1/2: target of rapamycin complex 1/2
UPR: unfolded protein response
WT: wildtype

## Supporting information

**Figure S1 - related to Figure 1.**
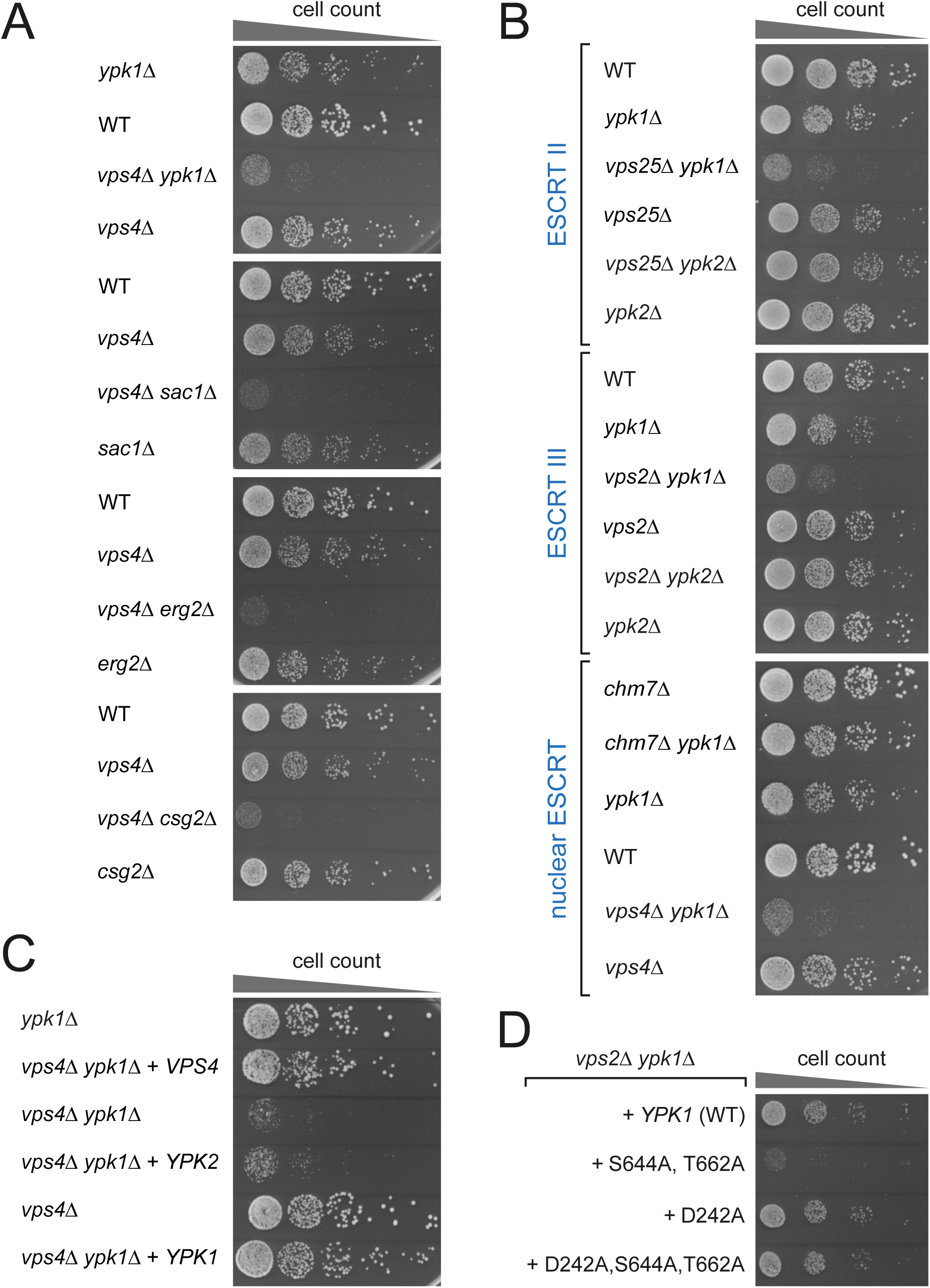
**(A, B)** Equal amounts of WT cells and indicated single or double mutants in serial dilutions were incubated on auxotrophic selection medium agar plates at 26°C. **(C)** Equal amounts of the indicated single or double mutants complemented with the indicated plasmids (or empty vectors for isogenic controls) in serial dilutions were incubated on auxotrophic selection medium agar plates at 26°C. **(D)** Equal amounts of *vps2Δ ypk1*Δ double mutants expressing the indicated *YPK1* mutant plasmids in serial dilutions were incubated on auxotrophic selection medium agar plates at 26°C.

**Figure S3 - related to Figure 3.**
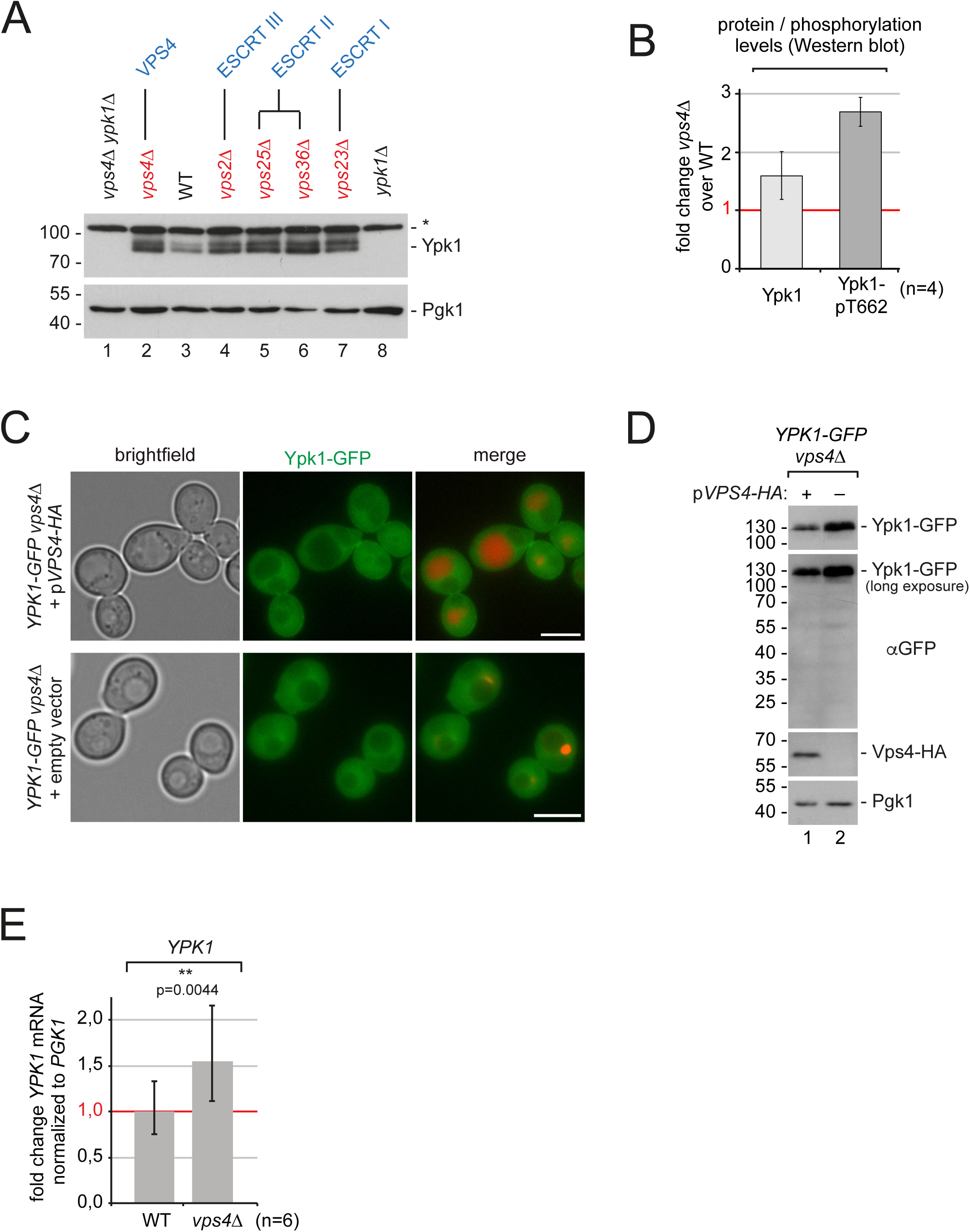
**(A)** SDS-PAGE and Western blot analysis with the indicated antibodies of total yeast lysates from WT and the indicated mutants. **(B)** Densitometric quantification of Ypk1 protein levels and Ypk1 pT662 phosphorylation levels in WT cells and *vps4*Δ mutants from Western blot experiments normalized to Pgk1 loading control. Data are presented as mean fold change in *vps4*Δ cells compared to WT levels ±standard deviation from 4 independent experiments. **(C)** Epifluorescence and phase contrast microscopy of living WT and *vps4*Δ cells expressing Ypk1-GFP (green) and the MVB cargo mCherry-CPS (red). Ypk1-GFP does not localize to the vacuole in WT or the class E compartment in ESCRT mutant cells grown under standard conditions. Scale bars 5µm. **(D)** SDS-PAGE and Western blot analysis with the indicated antibodies of total yeast lysates from WT and *vps4*Δ cells expressing *YPK1-GFP*. Only full-length Ypk1-GFP protein was observed in WT cells and ESCRT mutants. The typical free GFP fragment at 25 kD, which would be indicative of vacuolar proteolysis, was not detected. **(E)** Quantification of *YPK1* mRNA normalized to stable *PGK1* mRNA from WT cells and *vps4*Δ mutants by qPCR (n=6). Data are presented as mean fold change from WT ±standard deviation. Statistical significance was assessed by Student’s t-test.

**Figure S4 - related to Figure 4.**
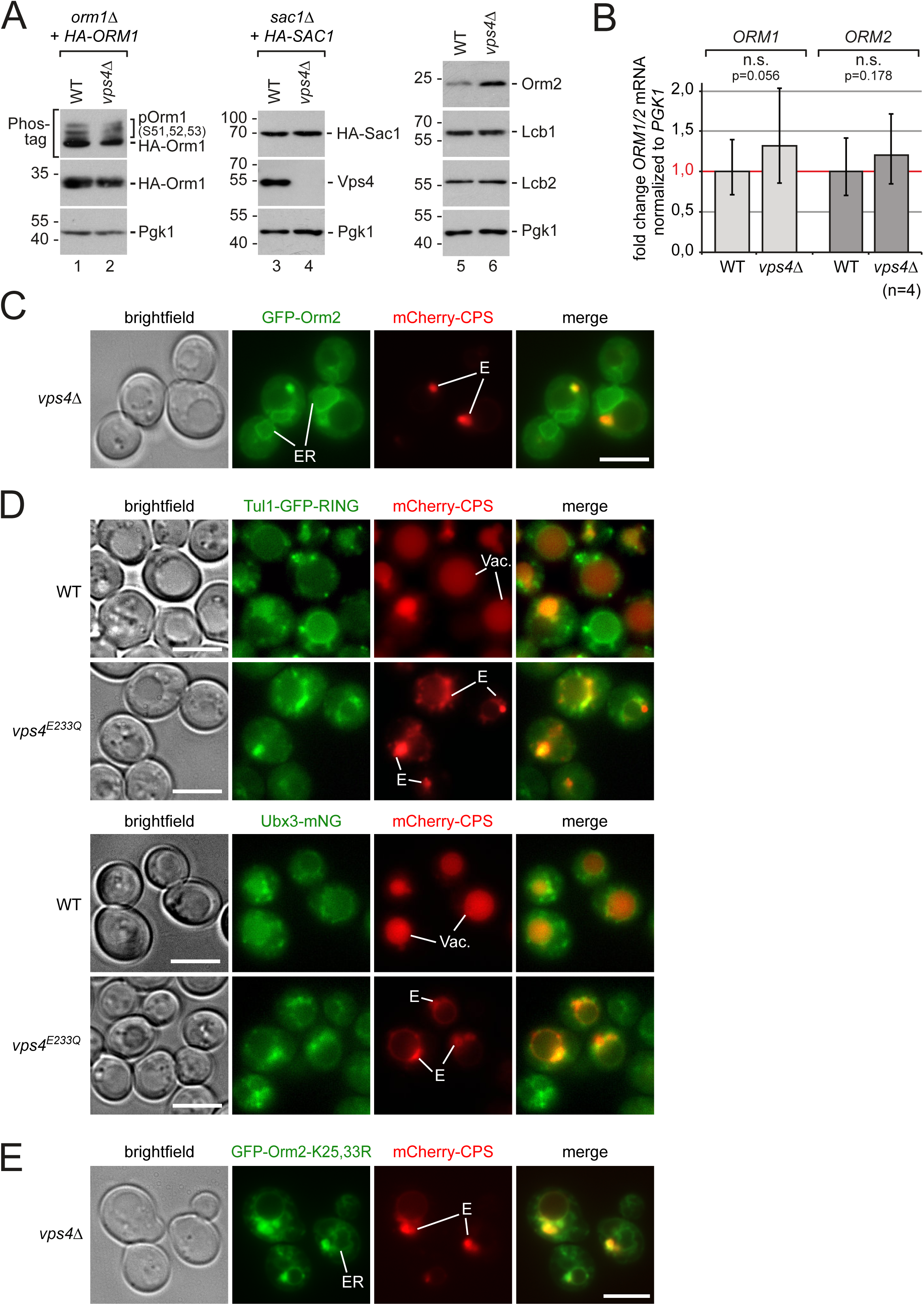
**(A)** SDS-PAGE and Western blot analysis with the indicated antibodies of total yeast lysates from the indicated WT or *vps4*Δ strains. **(B)** Quantification of *ORM1* and *ORM2* mRNA normalized to stable *PGK1* mRNA from WT cells and *vps4*Δ mutants by qPCR (n=4). Data are presented as mean fold change from WT ±standard deviation. Statistical significance was assessed by Student’s t-test. **(C)** Epifluorescence and phase contrast microscopy of living *vps4*Δ (*orm2*Δ) cells expressing GFP-Orm2 (green) and mCherry-CPS (red). ER, endoplasmic reticulum; E, class E compartment. **(D)** Epifluorescence and phase contrast microscopy of living *tul1*Δ cells expressing *TUL1-GFP-RING* (green) or WT cells expressing Ubx3-mNeonGreen (green) and mCherry-CPS (red), and either dominant-negative *vps4*E233Q or empty plasmid as indicated. Vac, vacuole; E, class E compartment. Scale bars 5µm. **(E)** Epifluorescence and phase contrast microscopy of living *vps4*Δ (*orm2*Δ) cells expressing GFP-Orm2-K25,33R (green) and mCherry-CPS (red). ER, endoplasmic reticulum; E, class E compartment.

**Figure S5 - related to Figure 5.**
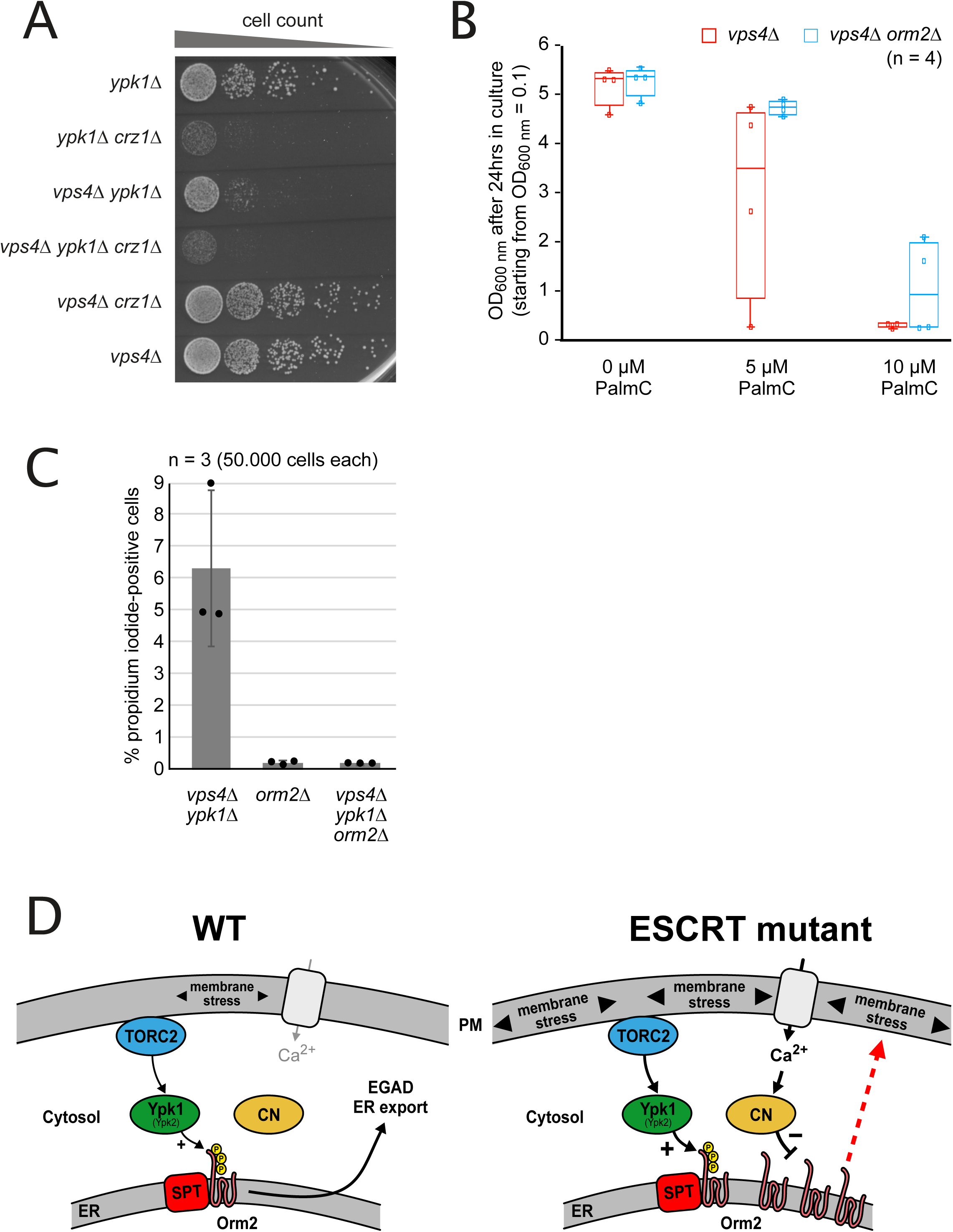
**(A)** Equal amounts of the indicated single, double or triple mutants in serial dilutions were incubated on a YPD agar plate at 26°C. **(B)** Growth of *vps4*Δ and *vps4*Δ *orm2*Δ cells in auxotrophic selection medium in presence of the indicated PalmC concentrations. Cells were inoculated to OD_600nm_ = 0.1 and grown for 24 hours in 4 independent experiments. The circles indicate the individual measurements. **(C)** The indicated strains were grown into mid-log phase in YPD medium and stained with propidium iodide for 10 minutes and analyzed by fluorescence activated cell sorting. Data are presented as mean ±standard deviation from 3 independent experiments. The data for the *vps4*Δ *ypk1*Δ strain is the same as in Fig. 2E and was plotted again for comparison. **(D)** Model for the post-translational regulation of Orm2 protein levels in WT cells and its deregulation in ESCRT mutants. PM, plasma membrane; ER, endoplasmic reticulum; TORC2, target of rapamycin complex 2; CN, calcineurin; SPT, serine-palmitoyl-coenzyme A transferase.

**Table S1:**
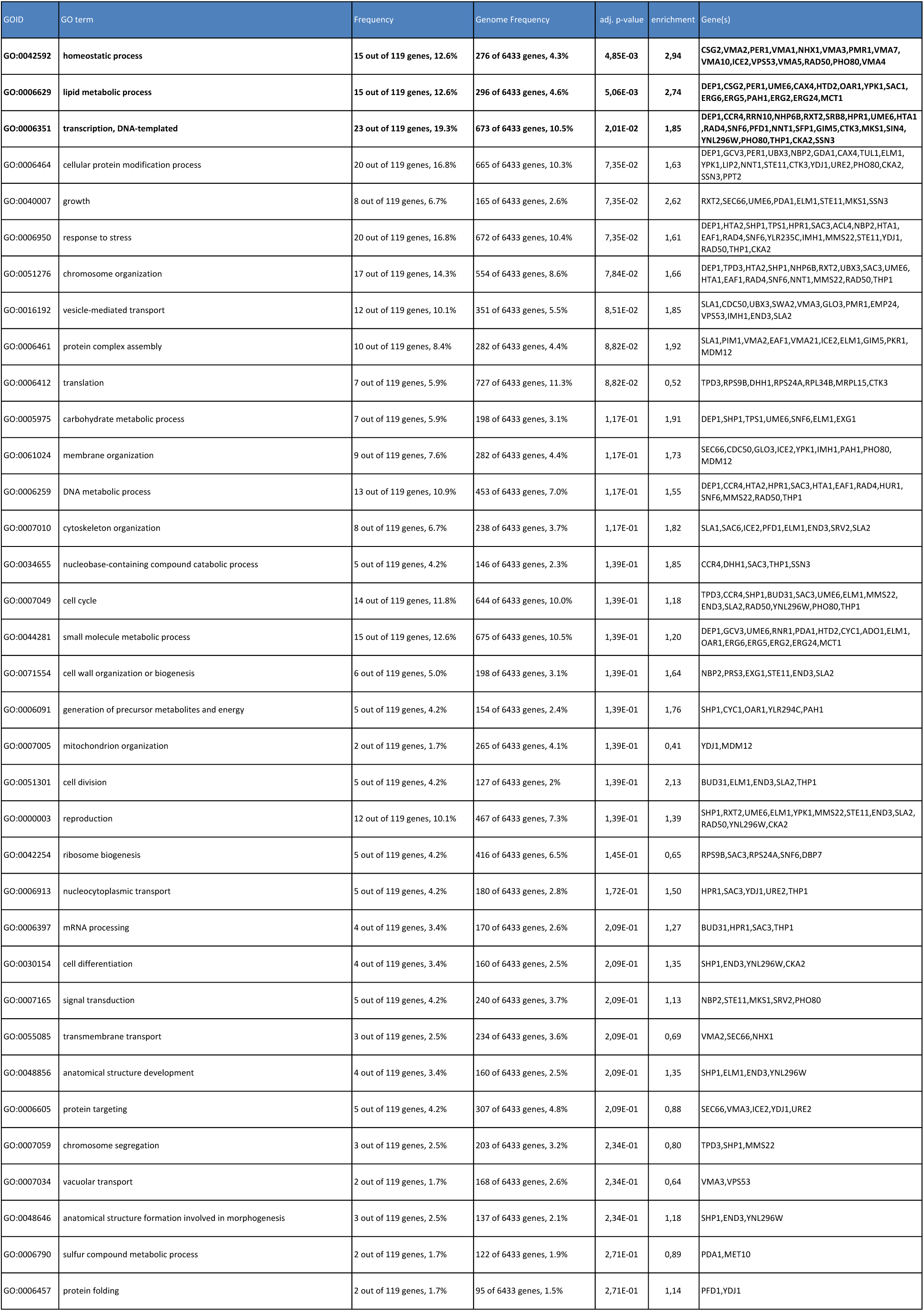
Gene ontology term ‘molecular processes’ enrichment analysis.

**Table S2:**
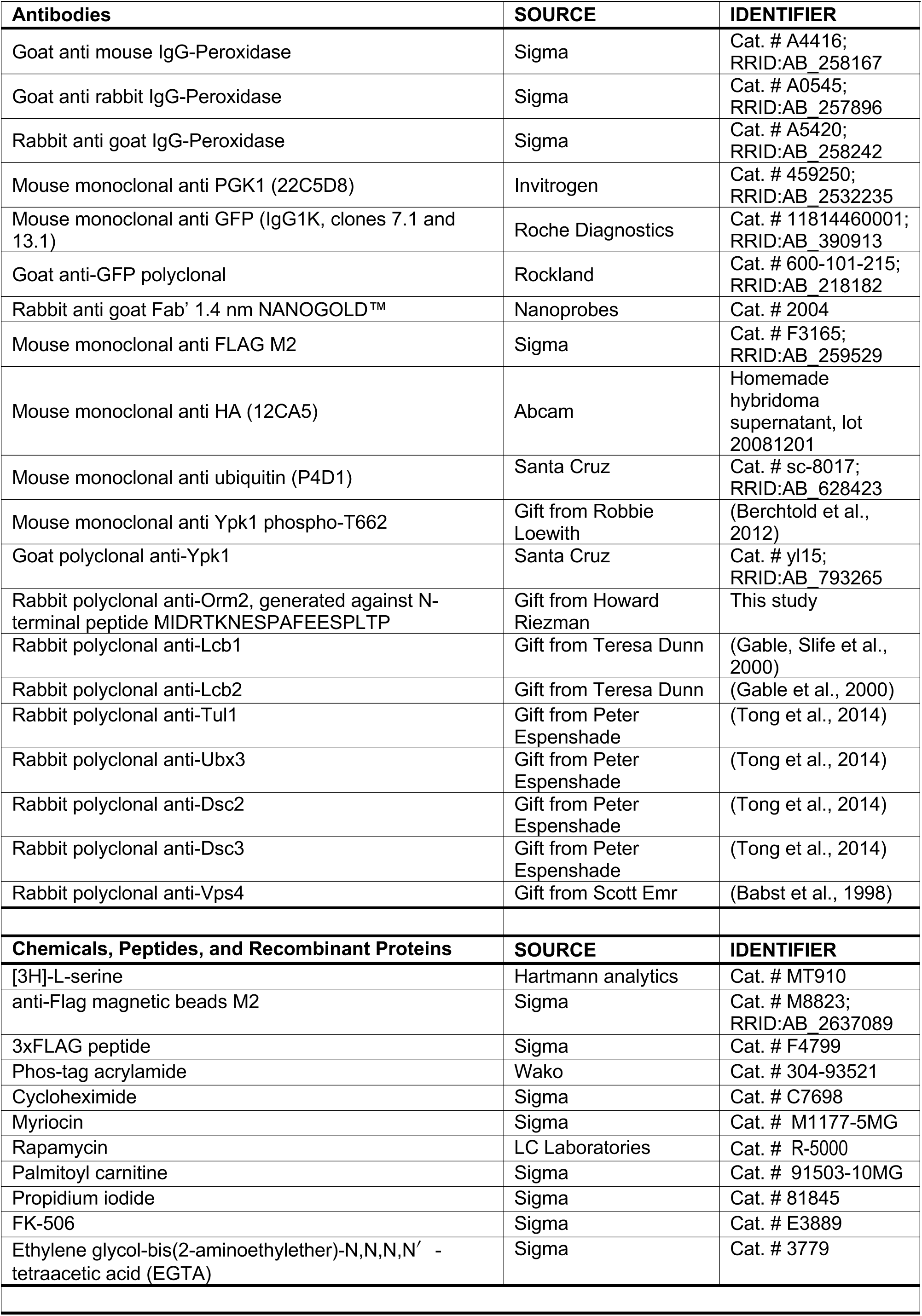
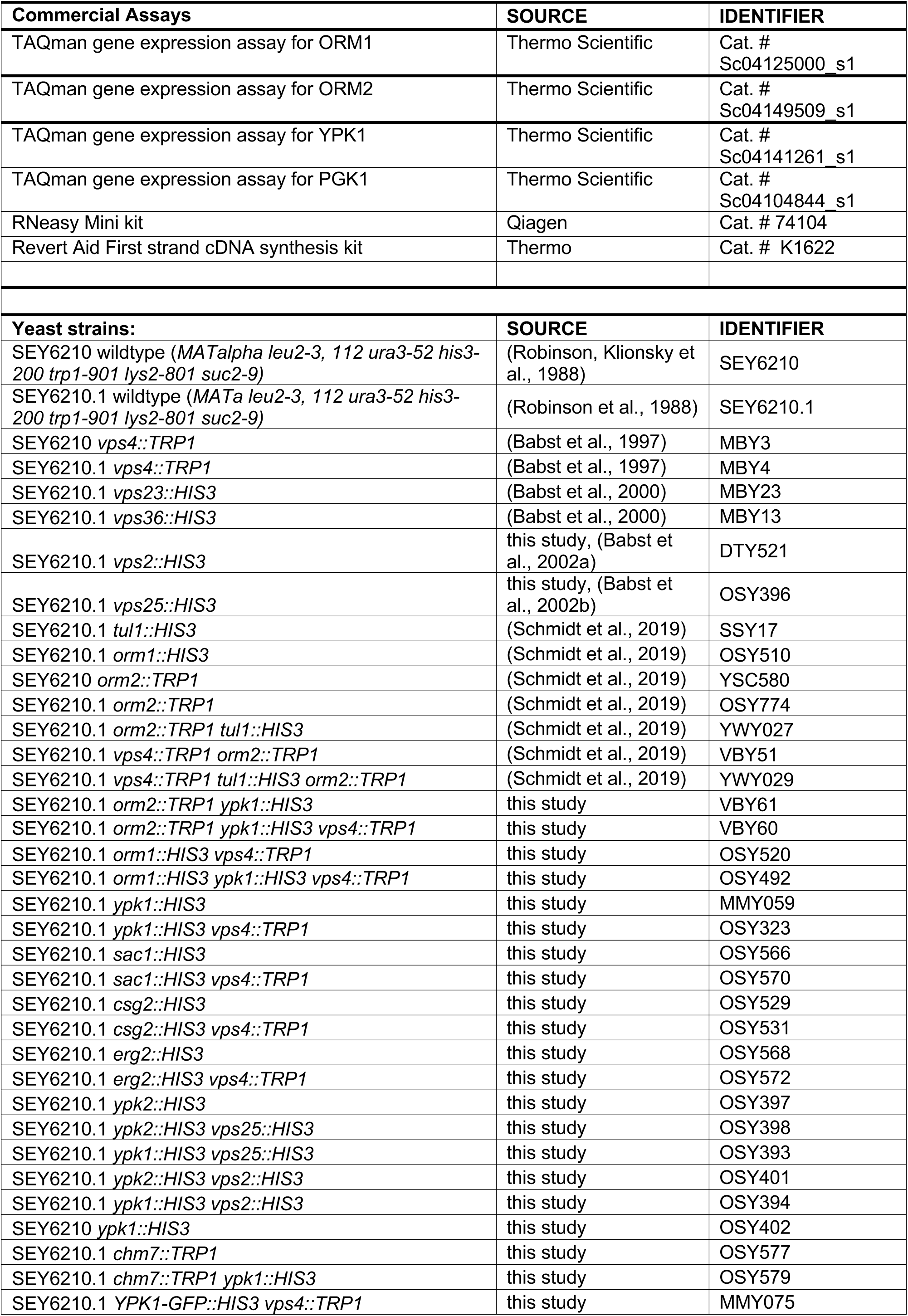
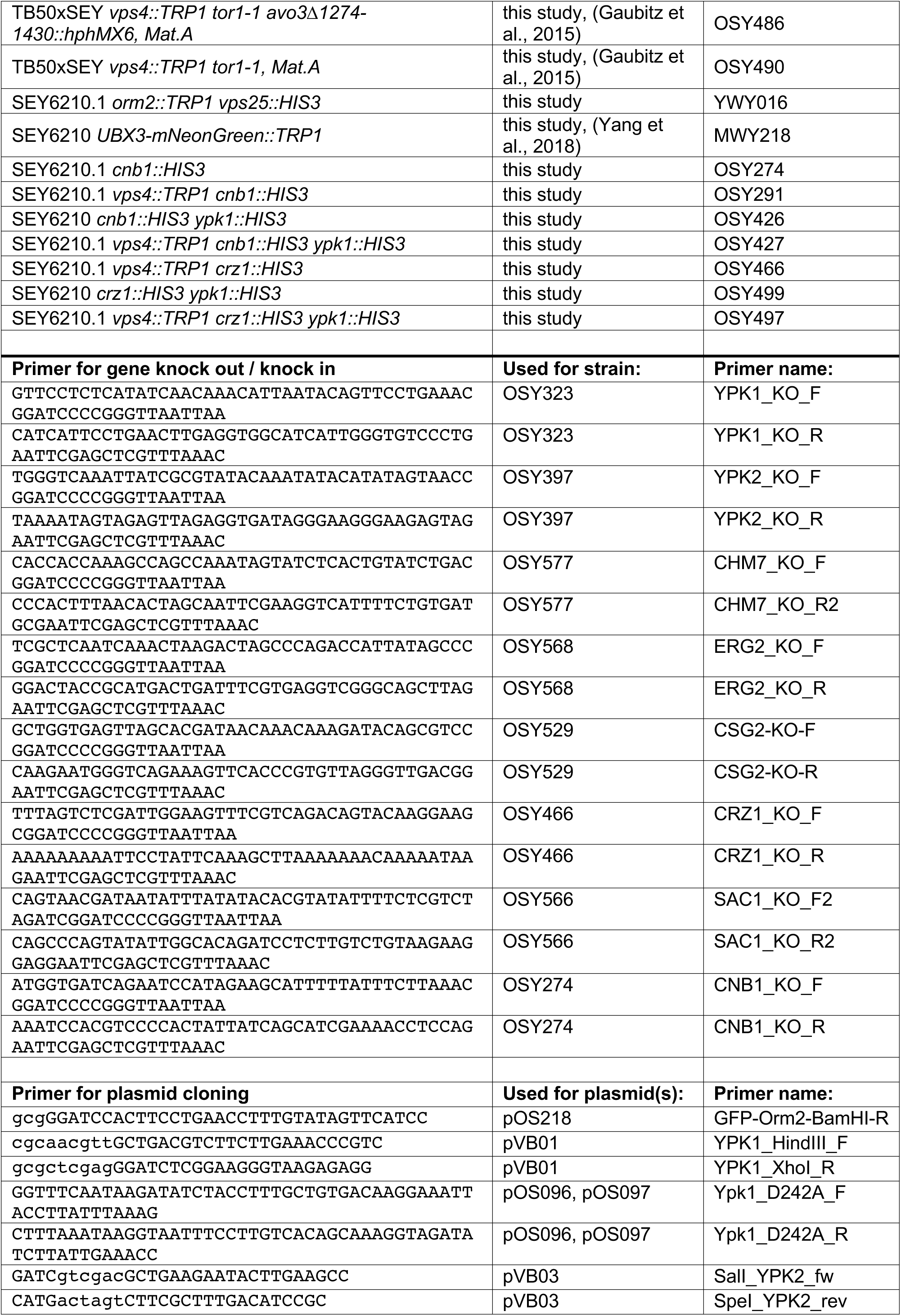
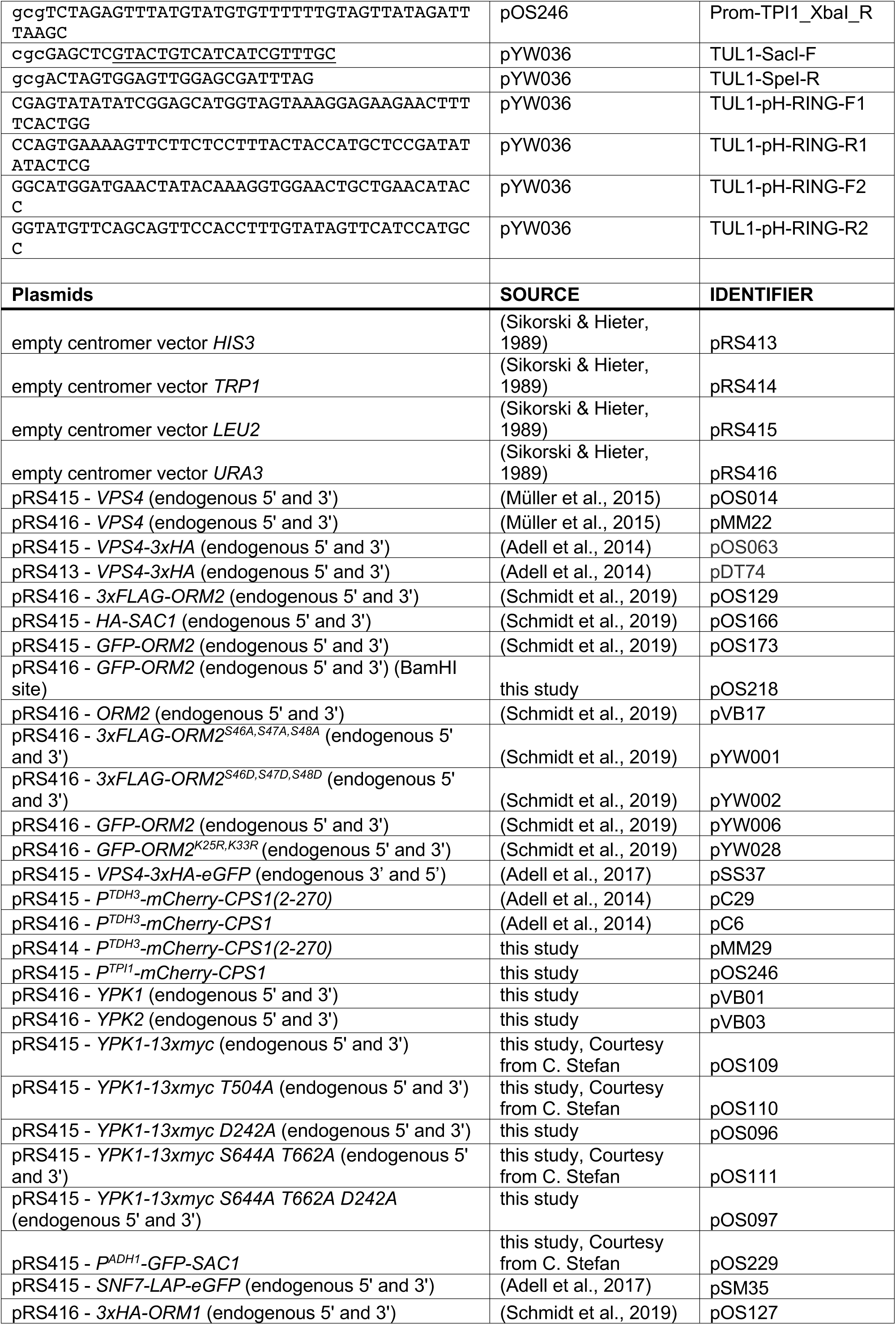
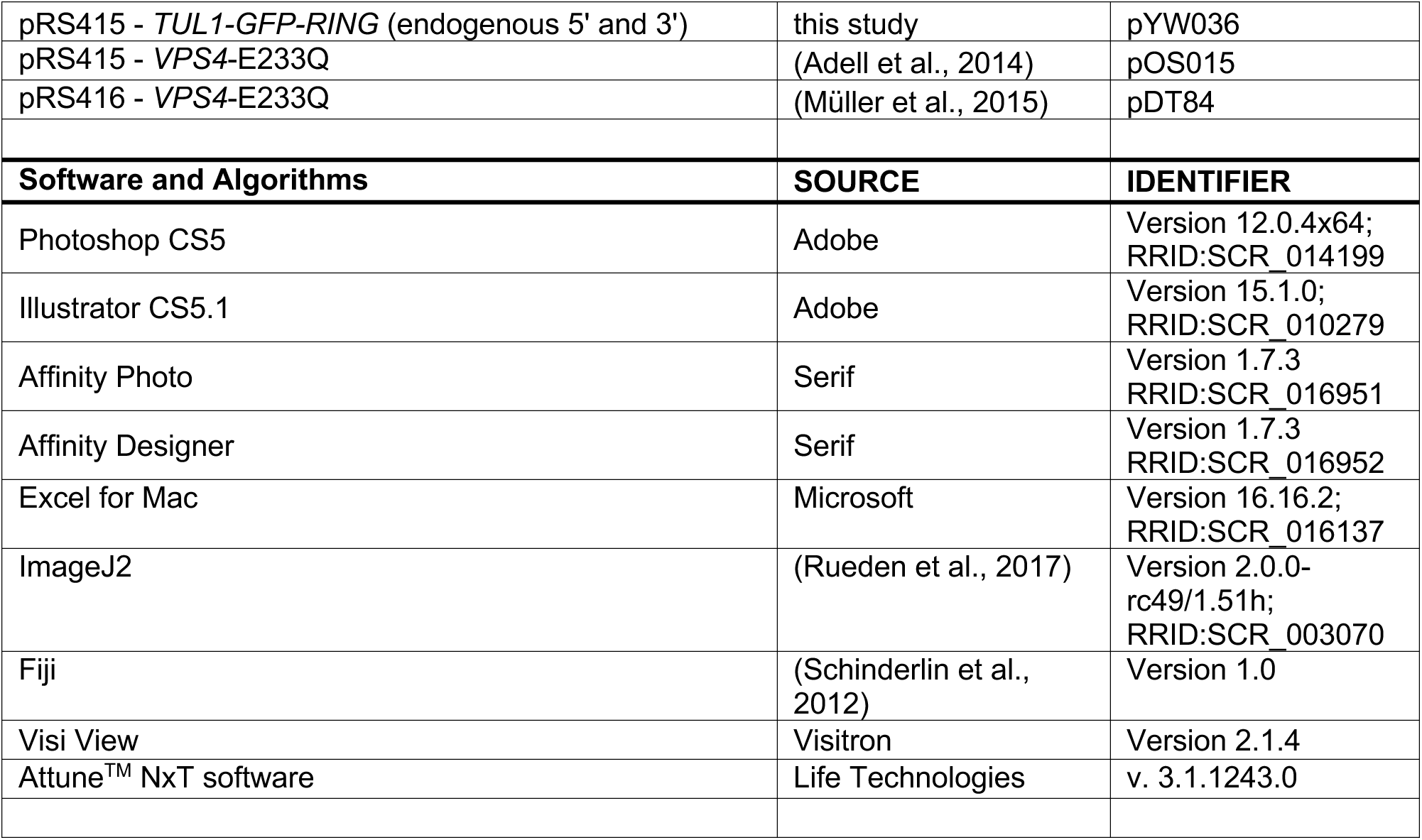
Yeast strains, plasmids and reagents.

## References

1. Ruggiano, A., Foresti, O., and Carvalho, P. (2014) Quality control: ER-associated degradation: protein quality control and beyond. J Cell Biol 204, 869–879

2. Wu, X., and Rapoport, T. A. (2018) Mechanistic insights into ER-associated protein degradation. Curr Opin Cell Biol 53, 22–28

3. Gil, G., Faust, J. R., Chin, D. J., Goldstein, J. L., and Brown, M. S. (1985) Membrane-bound domain of HMG CoA reductase is required for sterol-enhanced degradation of the enzyme. Cell 41, 249–258

4. Hampton, R. Y., Gardner, R. G., and Rine, J. (1996) Role of 26S proteasome and HRD genes in the degradation of 3-hydroxy-3-methylglutaryl-CoA reductase, an integral endoplasmic reticulum membrane protein. Mol Biol Cell 7, 2029–2044

5. Gill, S., Stevenson, J., Kristiana, I., and Brown, A. J. (2011) Cholesterol-dependent degradation of squalene monooxygenase, a control point in cholesterol synthesis beyond HMG-CoA reductase. Cell Metab 13, 260–273

6. Foresti, O., Ruggiano, A., Hannibal-Bach, H. K., Ejsing, C. S., and Carvalho, P. (2013) Sterol homeostasis requires regulated degradation of squalene monooxygenase by the ubiquitin ligase Doa10/Teb4. Elife 2, e00953

7. Strzyz, P. (2017) Unfolded protein response: Reacting to membrane stress. Nat Rev Mol Cell Biol 18, 471

8. Travers, K. J., Patil, C. K., Wodicka, L., Lockhart, D. J., Weissman, J. S., and Walter, P. (2000) Functional and genomic analyses reveal an essential coordination between the unfolded protein response and ER-associated degradation. Cell 101, 249–258

9. Schmidt, O., Weyer, Y., Baumann, V., Widerin, M. A., Eising, S., Angelova, M., Schleiffer, A., Kremser, L., Lindner, H., Peter, M., Frohlich, F., and Teis, D. (2019) Endosome and Golgi-associated degradation (EGAD) of membrane proteins regulates sphingolipid metabolism. EMBO J 38, e101433

10. Stewart, E. V., Nwosu, C. C., Tong, Z., Roguev, A., Cummins, T. D., Kim, D. U., Hayles, J., Park, H. O., Hoe, K. L., Powell, D. W., Krogan, N. J., and Espenshade, P. J. (2011) Yeast SREBP cleavage activation requires the Golgi Dsc E3 ligase complex. Mol Cell 42, 160–171

11. Reggiori, F., and Pelham, H. R. (2001) Sorting of proteins into multivesicular bodies: ubiquitin-dependent and -independent targeting. Embo J 20, 5176–5186

12. Li, M., Koshi, T., and Emr, S. D. (2015) Membrane-anchored ubiquitin ligase complex is required for the turnover of lysosomal membrane proteins. J Cell Biol 211, jcb.201505062-201505652

13. Dobzinski, N., Chuartzman, S. G., Kama, R., Schuldiner, M., and Gerst, J. E. (2015) Starvation-Dependent Regulation of Golgi Quality Control Links the TOR Signaling and Vacuolar Protein Sorting Pathways. Cell Rep

14. Tong, Z., Kim, M. S., Pandey, A., and Espenshade, P. J. (2014) Identification of candidate substrates for the Golgi Tul1 E3 ligase using quantitative diGly proteomics in yeast. Mol Cell Proteomics 13, 2871–2882

15. Breslow, D. K., Collins, S. R., Bodenmiller, B., Aebersold, R., Simons, K., Shevchenko, A., Ejsing, C. S., and Weissman, J. S. (2010) Orm family proteins mediate sphingolipid homeostasis. Nature 463, 1048–1053

16. Han, G., Gupta, S. D., Gable, K., Bacikova, D., Sengupta, N., Somashekarappa, N., Proia, R. L., Harmon, J. M., and Dunn, T. M. (2019) The ORMs interact with transmembrane domain 1 of Lcb1 and regulate serine palmitoyltransferase oligomerization, activity and localization. Biochim Biophys Acta Mol Cell Biol Lipids 1864, 245–259

17. Clarke, B. A., Majumder, S., Zhu, H., Lee, Y. T., Kono, M., Li, C., Khanna, C., Blain, H., Schwartz, R., Huso, V. L., Byrnes, C., Tuymetova, G., Dunn, T. M., Allende, M. L., and Proia, R. L. (2019) The Ormdl genes regulate the sphingolipid synthesis pathway to ensure proper myelination and neurologic function in mice. Elife 8

18. Moffatt, M. F., Kabesch, M., Liang, L., Dixon, A. L., Strachan, D., Heath, S., Depner, M., von Berg, A., Bufe, A., Rietschel, E., Heinzmann, A., Simma, B., Frischer, T., Willis-Owen, S. A., Wong, K. C., Illig, T., Vogelberg, C., Weiland, S. K., von Mutius, E., Abecasis, G. R., Farrall, M., Gut, I. G., Lathrop, G. M., and Cookson, W. O. (2007) Genetic variants regulating ORMDL3 expression contribute to the risk of childhood asthma. Nature 448, 470–473

19. Barrett, J. C., Hansoul, S., Nicolae, D. L., Cho, J. H., Duerr, R. H., Rioux, J. D., Brant, S. R., Silverberg, M. S., Taylor, K. D., Barmada, M. M., Bitton, A., Dassopoulos, T., Datta, L. W., Green, T., Griffiths, A. M., Kistner, E. O., Murtha, M. T., Regueiro, M. D., Rotter, J. I., Schumm, L. P., Steinhart, A. H., Targan, S. R., Xavier, R. J., Consortium, N. I. G., Libioulle, C., Sandor, C., Lathrop, M., Belaiche, J., Dewit, O., Gut, I., Heath, S., Laukens, D., Mni, M., Rutgeerts, P., Van Gossum, A., Zelenika, D., Franchimont, D., Hugot, J. P., de Vos, M., Vermeire, S., Louis, E., Belgian-French, I. B. D. C., Wellcome Trust Case Control, C., Cardon, L. R., Anderson, C. A., Drummond, H., Nimmo, E., Ahmad, T., Prescott, N. J., Onnie, C. M., Fisher, S. A., Marchini, J., Ghori, J., Bumpstead, S., Gwilliam, R., Tremelling, M., Deloukas, P., Mansfield, J., Jewell, D., Satsangi, J., Mathew, C. G., Parkes, M., Georges, M., and Daly, M. J. (2008) Genome-wide association defines more than 30 distinct susceptibility loci for Crohn’s disease. Nat Genet 40, 955–962

20. McGovern, D. P., Gardet, A., Torkvist, L., Goyette, P., Essers, J., Taylor, K. D., Neale, B. M., Ong, R. T., Lagace, C., Li, C., Green, T., Stevens, C. R., Beauchamp, C., Fleshner, P. R., Carlson, M., D’Amato, M., Halfvarson, J., Hibberd, M. L., Lordal, M., Padyukov, L., Andriulli, A., Colombo, E., Latiano, A., Palmieri, O., Bernard, E. J., Deslandres, C., Hommes, D. W., de Jong, D. J., Stokkers, P. C., Weersma, R. K., Consortium, N. I. G., Sharma, Y., Silverberg, M. S., Cho, J. H., Wu, J., Roeder, K., Brant, S. R., Schumm, L. P., Duerr, R. H., Dubinsky, M. C., Glazer, N. L., Haritunians, T., Ippoliti, A., Melmed, G. Y., Siscovick, D. S., Vasiliauskas, E. A., Targan, S. R., Annese, V., Wijmenga, C., Pettersson, S., Rotter, J. I., Xavier, R. J., Daly, M. J., Rioux, J. D., and Seielstad, M. (2010) Genome-wide association identifies multiple ulcerative colitis susceptibility loci. Nat Genet 42, 332–337

21. Barrett, J. C., Clayton, D. G., Concannon, P., Akolkar, B., Cooper, J. D., Erlich, H. A., Julier, C., Morahan, G., Nerup, J., Nierras, C., Plagnol, V., Pociot, F., Schuilenburg, H., Smyth, D. J., Stevens, H., Todd, J. A., Walker, N. M., Rich, S. S., and Type 1 Diabetes Genetics, C. (2009) Genome-wide association study and meta-analysis find that over 40 loci affect risk of type 1 diabetes. Nat Genet 41, 703–707

22. Roelants, F. M., Breslow, D. K., Muir, A., Weissman, J. S., and Thorner, J. (2011) Protein kinase Ypk1 phosphorylates regulatory proteins Orm1 and Orm2 to control sphingolipid homeostasis in Saccharomyces cerevisiae. Proc Natl Acad Sci U S A 108, 19222–19227

23. Kamada, Y., Fujioka, Y., Suzuki, N. N., Inagaki, F., Wullschleger, S., Loewith, R., Hall, M. N., and Ohsumi, Y. (2005) Tor2 directly phosphorylates the AGC kinase Ypk2 to regulate actin polarization. Mol Cell Biol 25, 7239–7248

24. Berchtold, D., Piccolis, M., Chiaruttini, N., Riezman, I., Riezman, H., Roux, A., Walther, T. C., and Loewith, R. (2012) Plasma membrane stress induces relocalization of Slm proteins and activation of TORC2 to promote sphingolipid synthesis. Nat Cell Biol 14, 542–547

25. Riggi, M., Niewola-Staszkowska, K., Chiaruttini, N., Colom, A., Kusmider, B., Mercier, V., Soleimanpour, S., Stahl, M., Matile, S., Roux, A., and Loewith, R. (2018) Decrease in plasma membrane tension triggers PtdIns(4,5)P2 phase separation to inactivate TORC2. Nat Cell Biol 20, 1043–1051

26. Aronova, S., Wedaman, K., Aronov, P. A., Fontes, K., Ramos, K., Hammock, B. D., and Powers, T. (2008) Regulation of ceramide biosynthesis by TOR complex 2. Cell Metab 7, 148–158

27. Berchtold, D., and Walther, T. C. (2009) TORC2 plasma membrane localization is essential for cell viability and restricted to a distinct domain. Mol Biol Cell 20, 1565–1575

28. Locke, M. N., and Thorner, J. (2019) Rab5 GTPases are required for optimal TORC2 function. J Cell Biol 218, 961–976

29. Niles, B. J., Mogri, H., Hill, A., Vlahakis, A., and Powers, T. (2012) Plasma membrane recruitment and activation of the AGC kinase Ypk1 is mediated by target of rapamycin complex 2 (TORC2) and its effector proteins Slm1 and Slm2. Proc Natl Acad Sci U S A 109, 1536–1541

30. Liao, H. C., and Chen, M. Y. (2012) Target of rapamycin complex 2 signals to downstream effector yeast protein kinase 2 (Ypk2) through adheres-voraciously-to-target-of-rapamycin-2 protein 1 (Avo1) in Saccharomyces cerevisiae. J Biol Chem 287, 6089–6099

31. Rispal, D., Eltschinger, S., Stahl, M., Vaga, S., Bodenmiller, B., Abraham, Y., Filipuzzi, I., Movva, N. R., Aebersold, R., Helliwell, S. B., and Loewith, R. (2015) Target of Rapamycin Complex 2 Regulates Actin Polarization and Endocytosis via Multiple Pathways. J Biol Chem 290, 14963–14978

32. Muir, A., Ramachandran, S., Roelants, F. M., Timmons, G., and Thorner, J. (2014) TORC2-dependent protein kinase Ypk1 phosphorylates ceramide synthase to stimulate synthesis of complex sphingolipids. Elife 3

33. Roelants, F. M., Baltz, A. G., Trott, A. E., Fereres, S., and Thorner, J. (2010) A protein kinase network regulates the function of aminophospholipid flippases. Proc Natl Acad Sci U S A 107, 34–39

34. Lee, Y. J., Jeschke, G. R., Roelants, F. M., Thorner, J., and Turk, B. E. (2012) Reciprocal phosphorylation of yeast glycerol-3-phosphate dehydrogenases in adaptation to distinct types of stress. Mol Cell Biol 32, 4705–4717

35. Muir, A., Roelants, F. M., Timmons, G., Leskoske, K. L., and Thorner, J. (2015) Down-regulation of TORC2-Ypk1 signaling promotes MAPK-independent survival under hyperosmotic stress. Elife 4

36. Roelants, F. M., Leskoske, K. L., Pedersen, R. T. A., Muir, A., Liu, J. M., Finnigan, G. C., and Thorner, J. (2017) TOR Complex 2-Regulated Protein Kinase Fpk1 Stimulates Endocytosis via Inhibition of Ark1/Prk1-Related Protein Kinase Akl1 in Saccharomyces cerevisiae. Mol Cell Biol 37

37. Hatakeyama, R., Kono, K., and Yoshida, S. (2017) Ypk1 and Ypk2 kinases maintain Rho1 at the plasma membrane by flippase-dependent lipid remodeling after membrane stresses. J Cell Sci 130, 1169–1178

38. Alvaro, C. G., Aindow, A., and Thorner, J. (2016) Differential Phosphorylation Provides a Switch to Control How alpha-Arrestin Rod1 Down-regulates Mating Pheromone Response in Saccharomyces cerevisiae. Genetics 203, 299–317

39. Chen, P., Lee, K. S., and Levin, D. E. (1993) A pair of putative protein kinase genes (YPK1 and YPK2) is required for cell growth in Saccharomyces cerevisiae. Mol Gen Genet 236, 443–447

40. Schöneberg, J., Lee, I.-H., Iwasa, J. H., and Hurley, J. H. (2017) Reverse-topology membrane scission by the ESCRT proteins. Nature reviews Molecular cell biology 18, 5–17

41. Vietri, M., Radulovic, M., and Stenmark, H. (2019) The many functions of ESCRTs. Nat Rev Mol Cell Biol

42. Migliano, S. M., and Teis, D. (2018) ESCRT and Membrane Protein Ubiquitination. Prog Mol Subcell Biol 57, 107–135

43. Odorizzi, G., Babst, M., and Emr, S. D. (1998) Fab1p PtdIns(3)P 5-kinase function essential for protein sorting in the multivesicular body. Cell 95, 847–858

44. Adell, M. A. Y., Migliano, S. M., Upadhyayula, S., Bykov, Y. S., Sprenger, S., Pakdel, M., Vogel, G. F., Jih, G., Skillern, W., Behrouzi, R., Babst, M., Schmidt, O., Hess, M. W., Briggs, J. A., Kirchhausen, T., and Teis, D. (2017) Recruitment dynamics of ESCRT-III and Vps4 to endosomes and implications for reverse membrane budding. Elife 6

45. Zhu, L., Jorgensen, J. R., Li, M., Chuang, Y. S., and Emr, S. D. (2017) ESCRTs function directly on the lysosome membrane to downregulate ubiquitinated lysosomal membrane proteins. Elife 6

46. Yang, X., Zhang, W., Wen, X., Bulinski, P. J., Chomchai, D. A., Arines, F. M., Liu, Y. Y., Sprenger, S., Teis, D., Klionsky, D. J., and Li, M. (2020) TORC1 regulates vacuole membrane composition through ubiquitin- and ESCRT-dependent microautophagy. J Cell Biol 219

47. Hatakeyama, R., Peli-Gulli, M. P., Hu, Z., Jaquenoud, M., Garcia Osuna, G. M., Sardu, A., Dengjel, J., and De Virgilio, C. (2019) Spatially Distinct Pools of TORC1 Balance Protein Homeostasis. Mol Cell 73, 325–338 e328

48. Raymond, C. K., Howald-Stevenson, I., Vater, C. A., and Stevens, T. H. (1992) Morphological classification of the yeast vacuolar protein sorting mutants: evidence for a prevacuolar compartment in class E vps mutants. Mol Biol Cell 3, 1389–1402

49. Gatta, A. T., and Carlton, J. G. (2019) The ESCRT-machinery: closing holes and expanding roles. Curr Opin Cell Biol 59, 121–132

50. Radulovic, M., Schink, K. O., Wenzel, E. M., Nahse, V., Bongiovanni, A., Lafont, F., and Stenmark, H. (2018) ESCRT-mediated lysosome repair precedes lysophagy and promotes cell survival. EMBO J 37

51. Vietri, M., Schink, K. O., Campsteijn, C., Wegner, C. S., Schultz, S. W., Christ, L., Thoresen, S. B., Brech, A., Raiborg, C., and Stenmark, H. (2015) Spastin and ESCRT-III coordinate mitotic spindle disassembly and nuclear envelope sealing. Nature 522, 231–235

52. Olmos, Y., Hodgson, L., Mantell, J., Verkade, P., and Carlton, J. G. (2015) ESCRT-III controls nuclear envelope reformation. Nature 522, 236–239

53. Webster, B. M., Colombi, P., Jager, J., and Lusk, C. P. (2014) Surveillance of nuclear pore complex assembly by ESCRT-III/Vps4. Cell 159, 388–401

54. Skowyra, M. L., Schlesinger, P. H., Naismith, T. V., and Hanson, P. I. (2018) Triggered recruitment of ESCRT machinery promotes endolysosomal repair. Science 360

55. McCullough, J., Clippinger, A. K., Talledge, N., Skowyra, M. L., Saunders, M. G., Naismith, T. V., Colf, L. A., Afonine, P., Arthur, C., Sundquist, W. I., Hanson, P. I., and Frost, A. (2015) Structure and membrane remodeling activity of ESCRT-III helical polymers. Science

56. Jimenez, A. J., Maiuri, P., Lafaurie-Janvore, J., Divoux, S., Piel, M., and Perez, F. (2014) ESCRT machinery is required for plasma membrane repair. Science 343, 1247136

57. Raab, M., Gentili, M., de Belly, H., Thiam, H. R., Vargas, P., Jimenez, A. J., Lautenschlaeger, F., Voituriez, R., Lennon-Dumenil, A. M., Manel, N., and Piel, M. (2016) ESCRT III repairs nuclear envelope ruptures during cell migration to limit DNA damage and cell death. Science 352, 359–362

58. Denais, C. M., Gilbert, R. M., Isermann, P., McGregor, A. L., te Lindert, M., Weigelin, B., Davidson, P. M., Friedl, P., Wolf, K., and Lammerding, J. (2016) Nuclear envelope rupture and repair during cancer cell migration. Science 352, 353–358

59. Roelants, F. M., Leskoske, K. L., Martinez Marshall, M. N., Locke, M. N., and Thorner, J. (2017) The TORC2-Dependent Signaling Network in the Yeast Saccharomyces cerevisiae. Biomolecules 7

60. Omnus, D. J., Manford, A. G., Bader, J. M., Emr, S. D., and Stefan, C. J. (2016) Phosphoinositide kinase signaling controls ER-PM cross-talk. Mol Biol Cell 27, 1170–1180

61. Bauer, I., Brune, T., Preiss, R., and Köllinger, R. (2015) Evidence for a Non-endosomal Function of the Saccharomyces cerevisiae ESCRT-III Like Protein Chm7. Genetics 115. 178939

62. Roelants, F. M., Torrance, P. D., Bezman, N., and Thorner, J. (2002) Pkh1 and Pkh2 differentially phosphorylate and activate Ypk1 and Ykr2 and define protein kinase modules required for maintenance of cell wall integrity. Mol Biol Cell 13, 3005–3028

63. deHart, A. K., Schnell, J. D., Allen, D. A., and Hicke, L. (2002) The conserved Pkh-Ypk kinase cascade is required for endocytosis in yeast. J Cell Biol 156, 241–248

64. Gaubitz, C., Oliveira, T. M., Prouteau, M., Leitner, A., Karuppasamy, M., Konstantinidou, G., Rispal, D., Eltschinger, S., Robinson, G. C., Thore, S., Aebersold, R., Schaffitzel, C., and Loewith, R. (2015) Molecular Basis of the Rapamycin Insensitivity of Target Of Rapamycin Complex 2. Mol Cell 58, 977–988

65. Riggi, M., Bourgoint, C., Macchione, M., Matile, S., Loewith, R., and Roux, A. (2019) TORC2 controls endocytosis through plasma membrane tension. J Cell Biol 218, 2265–2276

66. Ruhl, S., Shkarina, K., Demarco, B., Heilig, R., Santos, J. C., and Broz, P. (2018) ESCRT-dependent membrane repair negatively regulates pyroptosis downstream of GSDMD activation. Science 362, 956–960

67. Gong, Y. N., Guy, C., Olauson, H., Becker, J. U., Yang, M., Fitzgerald, P., Linkermann, A., and Green, D. R. (2017) ESCRT-III Acts Downstream of MLKL to Regulate Necroptotic Cell Death and Its Consequences. Cell 169, 286–300 e216

68. Scheffer, L. L., Sreetama, S. C., Sharma, N., Medikayala, S., Brown, K. J., Defour, A., and Jaiswal, J. K. (2014) Mechanism of Ca(2)(+)-triggered ESCRT assembly and regulation of cell membrane repair. Nat Commun 5, 5646

69. Leskoske, K. L., Roelants, F. M., Martinez Marshall, M. N., Hill, J. M., and Thorner, J. (2017) The Stress-Sensing TORC2 Complex Activates Yeast AGC-Family Protein Kinase Ypk1 at Multiple Novel Sites. Genetics 207, 179–195

70. Shimobayashi, M., Takematsu, H., Eiho, K., Yamane, Y., and Kozutsumi, Y. (2010) Identification of Ypk1 as a novel selective substrate for nitrogen starvation-triggered proteolysis requiring autophagy system and endosomal sorting complex required for transport (ESCRT) machinery components. J Biol Chem 285, 36984–36994

71. Guillas, I., Kirchman, P. A., Chuard, R., Pfefferli, M., Jiang, J. C., Jazwinski, S. M., and Conzelmann, A. (2001) C26-CoA-dependent ceramide synthesis of Saccharomyces cerevisiae is operated by Lag1p and Lac1p. EMBO J 20, 2655–2665

72. Schorling, S., Vallee, B., Barz, W. P., Riezman, H., and Oesterhelt, D. (2001) Lag1p and Lac1p are essential for the Acyl-CoA-dependent ceramide synthase reaction in Saccharomyces cerevisae. Mol Biol Cell 12, 3417–3427

73. Yang, X., Arines, F. M., Zhang, W., and Li, M. (2018) Sorting of a multi-subunit ubiquitin ligase complex in the endolysosome system. Elife 7

74. Mulet, J. M., Martin, D. E., Loewith, R., and Hall, M. N. (2006) Mutual antagonism of target of rapamycin and calcineurin signaling. J Biol Chem 281, 33000–33007

75. Zhao, Y., Du, J., Zhao, G., and Jiang, L. (2013) Activation of calcineurin is mainly responsible for the calcium sensitivity of gene deletion mutations in the genome of budding yeast. Genomics 101, 49–56

76. Zhao, Y., Du, J., Xiong, B., Xu, H., and Jiang, L. (2013) ESCRT components regulate the expression of the ER/Golgi calcium pump gene PMR1 through the Rim101/Nrg1 pathway in budding yeast. J Mol Cell Biol 5, 336–344

77. Cyert, M. S., and Thorner, J. (1992) Regulatory subunit (CNB1 gene product) of yeast Ca2+/calmodulin-dependent phosphoprotein phosphatases is required for adaptation to pheromone. Mol Cell Biol 12, 3460–3469

78. Gururaj, C., Federman, R. S., and Chang, A. (2013) Orm proteins integrate multiple signals to maintain sphingolipid homeostasis. J Biol Chem 288, 20453–20463

79. Yoshimoto, H., Saltsman, K., Gasch, A. P., Li, H. X., Ogawa, N., Botstein, D., Brown, P. O., and Cyert, M. S. (2002) Genome-wide analysis of gene expression regulated by the calcineurin/Crz1p signaling pathway in Saccharomyces cerevisiae. J Biol Chem 277, 31079–31088

80. Obara, K., and Kihara, A. (2014) Signaling events of the Rim101 pathway occur at the plasma membrane in a ubiquitination-dependent manner. Mol Cell Biol 34, 3525–3534

81. Lafaurie-Janvore, J., Maiuri, P., Wang, I., Pinot, M., Manneville, J. B., Betz, T., Balland, M., and Piel, M. (2013) ESCRT-III assembly and cytokinetic abscission are induced by tension release in the intercellular bridge. Science 339, 1625–1629

82. [preprint] Mercier, V., Larios, J., Molinard, G., Goujon, A., Matile, S., Gruenberg, J., and Roux, A. (2019) ENDOSOMAL MEMBRANE TENSION CONTROLS ESCRT-III-DEPENDENT INTRA-LUMENAL VESICLE FORMATION. bioRxiv, 550483

83. Bieberich, E. (2018) Sphingolipids and lipid rafts: Novel concepts and methods of analysis. Chem Phys Lipids 216, 114–131

84. Guan, X. L., Souza, C. M., Pichler, H., Dewhurst, G., Schaad, O., Kajiwara, K., Wakabayashi, H., Ivanova, T., Castillon, G. A., Piccolis, M., Abe, F., Loewith, R., Funato, K., Wenk, M. R., and Riezman, H. (2009) Functional interactions between sphingolipids and sterols in biological membranes regulating cell physiology. Mol Biol Cell 20, 2083–2095

85. Castro, B. M., Silva, L. C., Fedorov, A., de Almeida, R. F., and Prieto, M. (2009) Cholesterol-rich fluid membranes solubilize ceramide domains: implications for the structure and dynamics of mammalian intracellular and plasma membranes. J Biol Chem 284, 22978–22987

86. Megyeri, M., Riezman, H., Schuldiner, M., and Futerman, A. H. (2016) Making Sense of the Yeast Sphingolipid Pathway. J Mol Biol 428, 4765–4775

87. Walther, T. C., Brickner, J. H., Aguilar, P. S., Bernales, S., Pantoja, C., and Walter, P. (2006) Eisosomes mark static sites of endocytosis. Nature 439, 998–1003

88. Moseley, J. B. (2018) Eisosomes. Curr Biol 28, R376–R378

89. Athanasopoulos, A., Andre, B., Sophianopoulou, V., and Gournas, C. (2019) Fungal plasma membrane domains. FEMS Microbiol Rev

90. Appadurai, D., Gay, L., Moharir, A., Lang, M. J., Duncan, M. C., Schmidt, O., Teis, D., Vu, T. N., Silva, M., Jorgensen, E. M., and Babst, M. (2019) Plasma membrane tension regulates eisosome structure and function. Mol Biol Cell, mbcE19040218

91. Johnson, S. S., Hanson, P. K., Manoharlal, R., Brice, S. E., Cowart, L. A., and Moye-Rowley, W. S. (2010) Regulation of yeast nutrient permease endocytosis by ATP-binding cassette transporters and a seven-transmembrane protein, RSB1. J Biol Chem 285, 35792–35802

92. Worgall, T. S., Veerappan, A., Sung, B., Kim, B. I., Weiner, E., Bholah, R., Silver, R. B., Jiang, X. C., and Worgall, S. (2013) Impaired sphingolipid synthesis in the respiratory tract induces airway hyperreactivity. Sci Transl Med 5, 186ra167

93. Davis, D., Kannan, M., and Wattenberg, B. (2018) Orm/ORMDL proteins: Gate guardians and master regulators. Adv Biol Regul 70, 3–18

94. Frohlich, F., Petit, C., Kory, N., Christiano, R., Hannibal-Bach, H. K., Graham, M., Liu, X., Ejsing, C. S., Farese, R. V., and Walther, T. C. (2015) The GARP complex is required for cellular sphingolipid homeostasis. Elife 4

95. Olson, D. K., Frohlich, F., Farese, R. V., Jr., and Walther, T. C. (2016) Taming the sphinx: Mechanisms of cellular sphingolipid homeostasis. Biochim Biophys Acta 1861, 784–792

96. Zhao, Y., Xiong, B., Xu, H., and Jiang, L. (2014) Expression of NYV1 encoding the negative regulator of Pmc1 is repressed by two transcriptional repressors, Nrg1 and Mig1. FEBS Lett 588, 3195–3201

97. Müller, M., Schmidt, O., Angelova, M., Faserl, K., Weys, S., Kremser, L., Pfaffenwimmer, T., Dalik, T., Kraft, C., Trajanoski, Z., Lindner, H., Teis, D., and Mizushima, N. (2015) The coordinated action of the MVB pathway and autophagy ensures cell survival during starvation. Elife 4, e07736

98. Beeler, T., Gable, K., Zhao, C., and Dunn, T. (1994) A novel protein, CSG2p, is required for Ca2+ regulation in Saccharomyces cerevisiae. J Biol Chem 269, 7279–7284

99. Mishra, R., van Drogen, F., Dechant, R., Oh, S., Jeon, N. L., Lee, S. S., and Peter, M. (2017) Protein kinase C and calcineurin cooperatively mediate cell survival under compressive mechanical stress. Proc Natl Acad Sci U S A 114, 13471–13476

100. Goldman, A., Roy, J., Bodenmiller, B., Wanka, S., Landry, C. R., Aebersold, R., and Cyert, M. S. (2014) The calcineurin signaling network evolves via conserved kinase-phosphatase modules that transcend substrate identity. Mol Cell 55, 422–435

101. Rodriguez, A., Roy, J., Martinez-Martinez, S., Lopez-Maderuelo, M. D., Nino-Moreno, P., Orti, L., Pantoja-Uceda, D., Pineda-Lucena, A., Cyert, M. S., and Redondo, J. M. (2009) A conserved docking surface on calcineurin mediates interaction with substrates and immunosuppressants. Mol Cell 33, 616–626

102. Sun, Y., Miao, Y., Yamane, Y., Zhang, C., Shokat, K. M., Takematsu, H., Kozutsumi, Y., and Drubin, D. G. (2012) Orm protein phosphoregulation mediates transient sphingolipid biosynthesis response to heat stress via the Pkh-Ypk and Cdc55-PP2A pathways. Mol Biol Cell 23, 2388–2398

103. Vlahakis, A., Graef, M., Nunnari, J., and Powers, T. (2014) TOR complex 2-Ypk1 signaling is an essential positive regulator of the general amino acid control response and autophagy. Proc Natl Acad Sci U S A 111, 10586–10591

104. Zhao, Y., Macgurn, J. A., Liu, M., and Emr, S. (2013) The ART-Rsp5 ubiquitin ligase network comprises a plasma membrane quality control system that protects yeast cells from proteotoxic stress. Elife 2, e00459

105. Sikorski, R. S., and Hieter, P. (1989) A system of shuttle vectors and yeast host strains designed for efficient manipulation of DNA in Saccharomyces cerevisiae. Genetics 122, 19–27

106. Harris, M. A., Clark, J., Ireland, A., Lomax, J., Ashburner, M., Foulger, R., Eilbeck, K., Lewis, S., Marshall, B., Mungall, C., Richter, J., Rubin, G. M., Blake, J. A., Bult, C., Dolan, M., Drabkin, H., Eppig, J. T., Hill, D. P., Ni, L., Ringwald, M., Balakrishnan, R., Cherry, J. M., Christie, K. R., Costanzo, M. C., Dwight, S. S., Engel, S., Fisk, D. G., Hirschman, J. E., Hong, E. L., Nash, R. S., Sethuraman, A., Theesfeld, C. L., Botstein, D., Dolinski, K., Feierbach, B., Berardini, T., Mundodi, S., Rhee, S. Y., Apweiler, R., Barrell, D., Camon, E., Dimmer, E., Lee, V., Chisholm, R., Gaudet, P., Kibbe, W., Kishore, R., Schwarz, E. M., Sternberg, P., Gwinn, M., Hannick, L., Wortman, J., Berriman, M., Wood, V., de la Cruz, N., Tonellato, P., Jaiswal, P., Seigfried, T., and White, R. (2004) The Gene Ontology (GO) database and informatics resource. Nucleic Acids Res 32, D258–261

107. Kushnirov, V. V. (2000) Rapid and reliable protein extraction from yeast. Yeast 16, 857–860

108. Schmidt, O., Weyer, Y., Fink, M. J., Muller, M., Weys, S., Bindreither, M., and Teis, D. (2017) Regulation of Rab5 isoforms by transcriptional and post-transcriptional mechanisms in yeast. FEBS Lett 591, 2803–2815

109. Livak, K. J., and Schmittgen, T. D. (2001) Analysis of relative gene expression data using real-time quantitative PCR and the 2(-Delta Delta C(T)) Method. Methods 25, 402–408

110. van Donselaar, E., Posthuma, G., Zeuschner, D., Humbel, B. M., and Slot, J. W. (2007) Immunogold labeling of cryosections from high-pressure frozen cells. Traffic 8, 471–485

111. Tabuchi, M., Audhya, A., Parsons, A. B., Boone, C., and Emr, S. D. (2006) The phosphatidylinositol 4,5-biphosphate and TORC2 binding proteins Slm1 and Slm2 function in sphingolipid regulation. Mol Cell Biol 26, 5861–5875

112. Ejsing, C. S., Sampaio, J. L., Surendranath, V., Duchoslav, E., Ekroos, K., Klemm, R. W., Simons, K., and Shevchenko, A. (2009) Global analysis of the yeast lipidome by quantitative shotgun mass spectrometry. Proc Natl Acad Sci U S A 106, 2136–2141

113. Rueden, C. T., Schindelin, J., Hiner, M. C., DeZonia, B. E., Walter, A. E., Arena, E. T., and Eliceiri, K. W. (2017) ImageJ2: ImageJ for the next generation of scientific image data. BMC Bioinformatics 18, 529

